# Automatic Quality Control and Error Correction in MRI linear registration via a Residual Parameter Prediction Network for T1w MRI

**DOI:** 10.64898/2026.09.10.750693

**Authors:** Zhaojin Chen, Roqaie Moqadam, Amelie Metz, Walter Adame Gonzalez, the Alzheimer’s Disease Neuroimaging Initiative (ADNI), the Consortium for the early identification of Alzheimer’s Disease–Quebec (CIMA-Q), The PREVENT-AD Research Group, Yashar Zeighami, Mahsa Dadar

## Abstract

Errors in linear registration can propagate to downstream nonlinear registration and bias volumetric estimations, deformation-based morphometry (DBM) and voxel-based morphometry (VBM) analyses. Subtle linear registration errors are particularly challenging as they are difficult to detect and may not result in obvious failures in nonlinear registration but still affect downstream results. Therefore, accurate identification and correction of these errors are critical. In this study, we present the Residual Affine COefficient Optimization Network (RACOON), a framework designed to identify and correct linear registration errors in T1w MRI scans registered to the MNI-ICBM152 space. RACOON’s correction module achieved a residual misalignment RMSE of 0.778 mm on synthetic dataset, comparable to the variability observed among repeated QC-passed registrations using the same pipeline. For the classification module, RACOON achieved a balanced accuracy of 76.8% and a precision of 74.4%, outperforming existing state-of-the-art methods. RACOON is open source and publicly available at https://github.com/ZhaojinChen/RACOON.

## 1. Introduction

Linear registration establishes a global alignment between an individual MRI scan and a second reference scan, commonly a template. Linear registration is one of the most commonly used initial steps in all image processing pipelines, and its inaccuracy would substantially impact many downstream derivatives. For example,the estimated scale factors from linear registration are commonly used for the estimation of total intracranial volume (Buckner et al., 2004; Dadar et al., 2025; Im et al., 2008). Prior to nonlinear registration, linear registration is also the first registration step in deformation-based morphometry (DBM) and voxel-based morphometry (VBM) analyses, allowing for voxel-level characterization and longitudinal tracking of brain atrophy (Baron et al., 2001; Manera et al., 2019). Previous studies have shown that registration errors can bias downstream analyses and alter the derived patterns of regional atrophy (Ceccarelli et al., 2012; Kim et al., 2015; Metz et al., 2025). Thus, all image processing studies should verify the accuracy of the linear registrations before proceeding with downstream analyses.

Several visual inspection protocols have been developed and validated for linear registration QC in recent years (Benhajali et al., 2020; Dadar et al., 2018; Fernandez-Lozano et al., 2024; Raamana et al., 2020). Most protocols compare the overlap between the template outline or specific landmarks and the transformed individual MRI scans. Although establishing consistent and replicable quality control procedures is vital, given the rapid growth of open-access MRI datasets from large-scale projects such as the UK Biobank and the time necessary to perform manual assessments (∼30 hours for ∼10K registrations (Dadar et al., 2018)), fully manual QC is becoming increasingly impractical (Markiewicz et al., 2021; Miller et al., 2016).

Most prior studies in the literature focus on assessing the overall registration quality by comparing the transformed segmentation labels with the template labels using overlap- or distance-based metrics (De Vos et al., 2019; Dubost et al., 2020; Hoffmann et al., 2021; Iglesias, 2023). However, large linear-registration errors can lead to failure of subsequent nonlinear registrations, while subtle linear-registration errors may be partially compensated by nonlinear registration but still bias deformation estimates. Thus, independent QC for linear registration is essential. Several automated methods have been proposed for linear registration QC in the literature. Mirhakimi et al. observed that MRI scans that failed QC showed substantially lower numerical stability than those that passed QC and proposed numerical stability defined as variability in computational results introduced by floating-point errors for a potential QC measure (Mirhakimi et al., 2025). Tummala and colleagues generated synthetic misaligned MRI scans and used similarity metrics, such as normalized cross-correlation, to distinguish misaligned from correctly aligned T1-weighted (T1w) and T2-weighted MRI scans (Tummala et al., 2021). However, because of individual anatomical variability and age-related atrophy or pathology, approaches using similarity-based cost functions have difficulty identifying subtle registration errors. de Senneville et al. proposed RegQCNET, a network that estimates the misalignment distance relative to a gold-standard (QC-passed) registration and identifies QC-failed MRI scans by applying a threshold to the estimated distance (de Senneville et al., 2020). Fonov and colleagues proposed a direct classification approach DARQ by fine-tuning a pretrained ResNet using the middle three slices of each MRI scan (V. S. Fonov et al., 2022). Thadikemalla et al. used the reconstruction error as the QC metric, assuming that the reconstruction error would increase as misalignment becomes more severe (Thadikemalla et al., 2024).

However, several important issues remain overlooked or underexplored in the aforementioned existing automated QC approaches. First, most existing QC models are developed on preprocessed MRI images, hindering their ability to generalize to different pipelines with inevitably different preprocessing methods applied. Second, many publicly available datasets deface MRI scans (e.g. CamCAN and UK Biobank) (Miller et al., 2016; Shafto et al., 2014), leading to reduced generalizability for many methods, as the potential effects of different defacing techniques have not been considered in existing registration QC models. Third, the classification performance of current models can be affected by intra- and inter-rater variability in manual QC labels, which may limit their direct application across laboratories with different QC criteria. In addition, subtle registration errors remain particularly challenging to identify, as most existing automated QC methods have primarily focused on relatively large registration errors. Lastly, images that fail QC are typically excluded from subsequent analyses. Previous studies have shown that failed cases often present with severe atrophy, increased white matter hyperintensity burden, or greater pathological burden (Dadar et al., 2018; Metz et al., 2025). Consequently, excluding these images may introduce bias in downstream analyses.

To address this limitation, we propose a Residual Affine COefficient Optimization Network (RACOON), a fast and robust framework that accurately identifies linear registration errors and, more importantly, corrects misalignment errors in QC-failed T1w images. Our proposed model can generalize across different scanners, field strengths, defacing methods, and preprocessing pipelines, while improving the identification of subtle linear registration errors compared with current automatic methods.

## 2. Methods

### 2.1 Datasets

To develop a robust and generalizable model, we aggregated and used T1w MRI scans from 13 publicly available datasets. These datasets included scans acquired at different image resolutions (Table 1), scanners (Siemens, GE, and Philips), and field strengths (1.5 T and 3.0 T). These data also cover diverse clinical populations, including healthy aging, frontotemporal dementia, Parkinson’s disease, and Alzheimer’s disease. Detailed descriptions of all datasets used in this study are provided in the Supplementary S1. Three subsets of the 13 datasets, referred to as Datasets A, B, and C, were constructed for different experiments, as detailed in Supplementary S2. Dataset A was used for the template-input ablation experiment and hyperparameter tuning, while Dataset B was used to evaluate the effect of training sample size. Dataset C consisted of equal numbers of real QC-passed and QC-failed scans with manual QC labels and was used for training and evaluating the classification module.

**Table 1.** MRI acquisition characteristics and number of scans included for each dataset.

| Datasets | Field strength | Scanner | Resolution [mm<sup>3</sup>] | N total scans | N QC-failed scans |
| --- | --- | --- | --- | --- | --- |
| UKBB | 3.0T | Siemens | 1.0×1.0×1.0 | 15242 | 2871 |
| ADNI | 1.5T, 3.0T | Siemens, GE, Philips | 1.2×1.2×1.2<br>1.2×1.0×1.0 | 388 | 48 |
| PPMI | 1.5T, 3.0T | Siemens, GE, Philips | 1.2×1.0×1.0<br>1.0×1.0×1.0 | 3133 | 365 |
| CamCAN | 3.0T | Siemens | 1.0×1.0×1.0 | 650 | 20 |
| HABS-Aging | 3.0T | Siemens | 1.0×1.0×1.0 | 278 | 10 |
| NACC | 1.5T, 3.0T | Siemens, GE, Philips | 1.2×1.0×1.0<br>1.0×1.0×1.0<br>1.0×1.5×1.0 | 2920 | 162 |
| NIFD | 3.0T | Siemens, GE | 1.2×1.0×1.0<br>1.0×1.0×1.0 | 339 | 3 |
| CCNA | 3.0T | Siemens, GE, Philips | 1.0×1.0×1.0 | 928 | 76 |
| CIMA-Q | 3.0T | Siemens, Philips | 1.0×1.0×1.0 | 287 | 76 |
| ALLFTD | 3.0T | Siemens, GE, Philips | 1.2×1.0×1.0<br>1.2×1.1×1.1<br>1.0×1.0×1.0 | 1837 | 14 |
| PrevenAD | 3.0T | Siemens | 1.0×1.0×1.0 | 355 | 28 |
| HCP Aging | 3.0T | Siemens | 0.8×0.8×0.8 | 1054 | 6 |
| HCP Adults | 3.0T | Siemens | 0.7×0.7×0.7 | 650 | 8 |

**Abbreviations:** UKBB: UK Biobank; ADNI: Alzheimer’s Disease Neuroimaging Initiative; PPMI: Parkinson’s Progression Markers Initiative; CamCAN: Cambridge Centre for Ageing and Neuroscience; HABS-Aging: Harvard Aging Brain Study–Aging; NACC: National Alzheimer’s Coordinating Center; NIFD: Frontotemporal Lobar Degeneration Neuroimaging Initiative; CCNA: Canadian Consortium on Neurodegeneration in Aging; CIMA-Q: Consortium for the Early Identification of Alzheimer’s Disease–Québec; ALLFTD: ARTFL/LEFFTDS Longitudinal Frontotemporal Lobar Degeneration; PrevenAD: Pre-symptomatic Evaluation of Experimental or Novel Treatments for Alzheimer’s Disease; HCP Aging: Human Connectome Project.
**N QC-failed:** Number of scans that failed visual quality control (QC).

**Table 2.** MAE of affine parameters and RMSE between repeated QC-passed registrations across ADNI, UKBB, and PPMI.

| Dataset | Number of Scans | RMSE (between 2 runs) | Axis | Rotation | Scale | Translation |
| --- | --- | --- | --- | --- | --- | --- |
| ADNI | 1948 | 0.786±0.531 | X | 0.329±0.385 | 0.005±0.005 | 0.139±0.120 |
|  |  |  | Y | 0.129±0.115 | 0.004±0.005 | 0.236±0.245 |
|  |  |  | Z | 0.101±0.096 | 0.008±0.008 | 0.265±0.258 |
| PPMI | 1922 | 0.357±0.389 | X | 0.166±0.243 | 0.002±0.002 | 0.068±0.118 |
|  |  |  | Y | 0.062±0.067 | 0.002±0.003 | 0.104±0.194 |
|  |  |  | Z | 0.047±0.052 | 0.003±0.004 | 0.110±0.265 |
| UKBB | 248 | 0.281±0.176 | X | 0.116±0.109 | 0.001±0.001 | 0.055±0.043 |
|  |  |  | Y | 0.044±0.044 | 0.001±0.001 | 0.104±0.101 |
|  |  |  | Z | 0.038±0.038 | 0.002±0.003 | 0.111±0.107 |

### 2.2 Manual Quality Control Criteria

Each T1-weighted MRI scan registered in ICBM space was visually inspected by expert raters and assigned a binary label (Pass or Fail) indicating whether the image was well aligned with the template using the estimated affine transformation by Advanced Normalization Tools (ANTs)(Avants et al., 2009). Quality control was performed by visually assessing the alignment between the template outline and the individual MRI scan according to the following criteria:

1. Sagittal and coronal views were first assessed to identify potential scaling errors.
2. If these views were well aligned, axial slices were assessed for rotation errors in the axial plane, particularly misalignment in the frontal lobe.
3. A scan is considered a failure when the mismatch is visible across multiple slices, indicating a scaling or rotation error rather than normal anatomical variation.
4. The brain axis in the axial view is also examined to ensure that it is aligned with the template (although this type of failure is uncommon).
5. Finally, the central gyrus is examined to ensure its alignment with the template.
6. Ventricular alignment was not used as a QC landmark because ventricular enlargement is common in aging and neurodegenerative disorders.

Linear registration errors are most commonly due to scaling and rotation misestimations. Examples of QC-passed and QC-failed MRI scans are shown in Fig. 1. For QC-passed scans, the template outlines closely match the brain across all 60 displayed slices, as illustrated in Fig. 1A and Fig. 1C. The QC-failed scan in Fig. 1B shows an underestimated scaling factor along the z-axis, resulting in a clear mismatch between the brain and template outlines in both the sagittal and coronal views (indicated by red arrows). Fig. 1D shows a rotation error, with visible misalignment of the frontal lobe in the axial view and the temporal lobe in the sagittal view (indicated by red arrows). In contrast, although Fig. 1C also shows some mismatch in the frontal lobe, the temporal lobe remains well aligned with the template outline. This pattern suggests that the frontal mismatch is more likely related to atrophy than to a registration error.

**Fig. 1.**
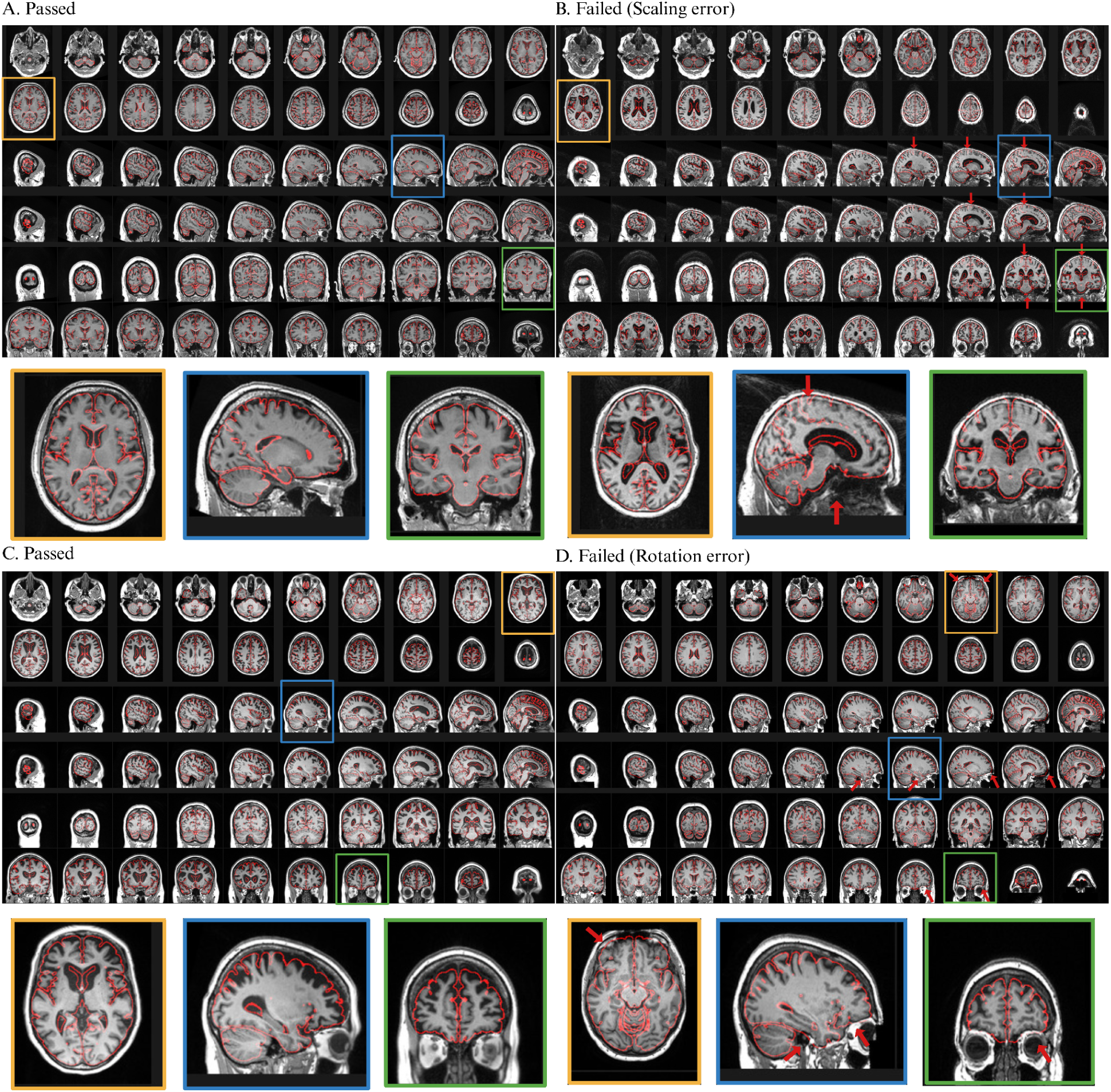
Examples of QC-passed and QC-failed MRI scans. Twenty slices were displayed in each view (i.e., axial, coronal, and sagittal), resulting in a total of 60 slices for each scan. The red contours represent the template outline, and the red arrows in B and D indicate regions of registration misalignment. Selected slices are enlarged and shown below the corresponding images for better visualization.

### 2.3 RACOON framework

The overall framework of RACOON is shown in Fig. 2. RACOON consists of three modules: (A) residual parameter prediction, (B) classification, and (C) correction. The three modules were designed based on three assumptions:

1. QC-passed registrations require no further adjustment of their affine parameters. Therefore, synthetically misaligned MRIs can be generated from QC-passed scans by applying extra affine parameters, allowing a model to learn to predict these parameters from the misaligned images.
2. QC-passed and QC-failed registrations have different residual affine parameters, which can be used to distinguish between the two groups. More specifically, the visual QC pass or fail status can be predicted based on the combined set of the residual affined parameters. This module can be further adapted for different levels of sensitivity.
3. The residual affine parameters estimated from QC-failed registrations can be used to correct their misalignment and recover a QC-passed alignment.

**Fig. 2.**
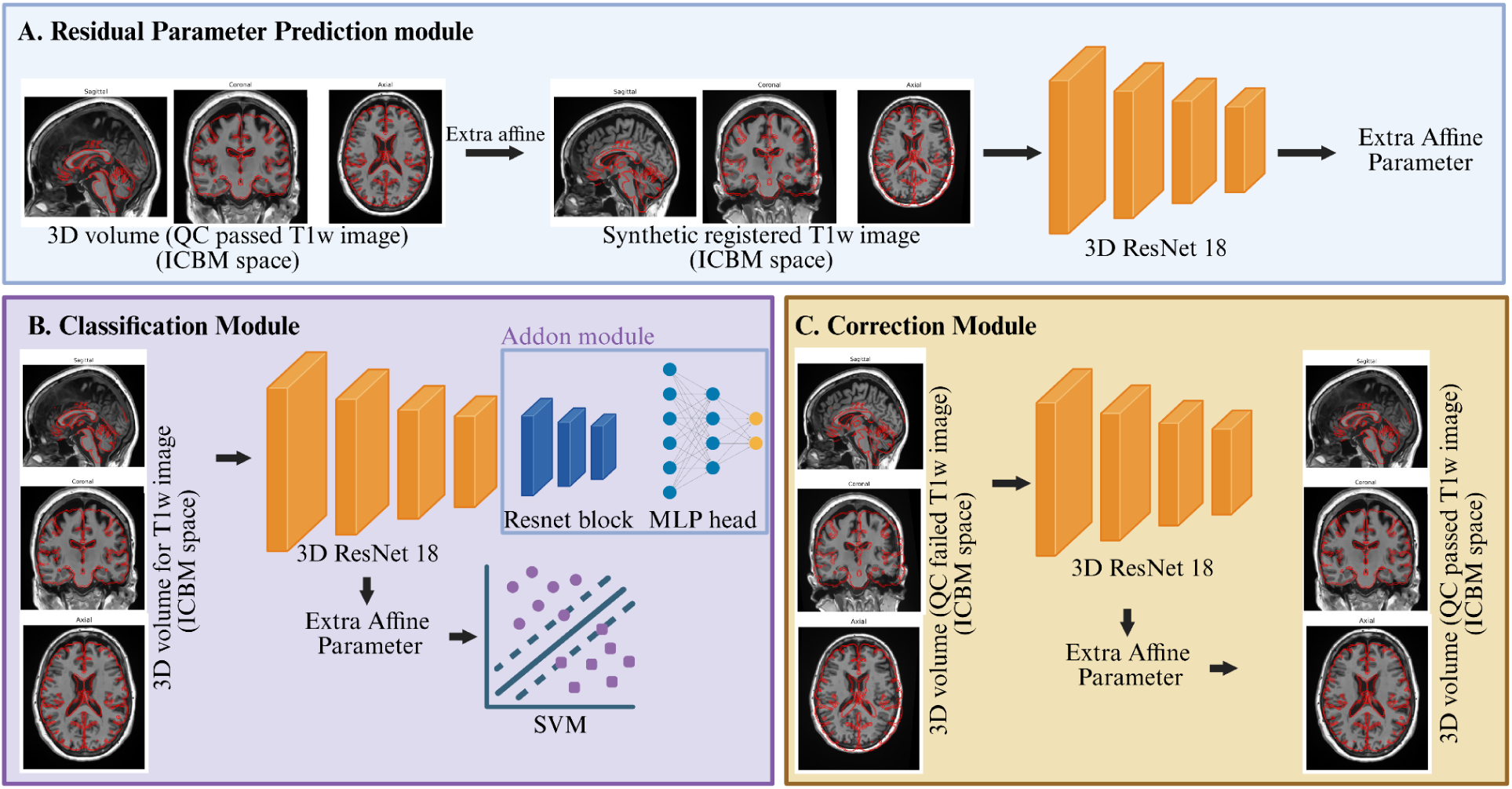
RACOON Framework. **(A)** An extra affine matrix is applied to a QC-passed T1w MRI scan to get the synthetic misaligned MRIs, and a 3D ResNet18 (yellow blocks) is trained to predict the applied affine parameters. **(B)** Two classification approaches are illustrated. The first extends the frozen ResNet18 (yellow blocks) with additional residual blocks (blue blocks) and an MLP head for binary QC classification. The second uses the predicted affine parameters as input to an SVM classifier. **(C)** For QC-failed scans, the affine parameters predicted by the 3D ResNet18 are used to correct the registration error and improve alignment.

#### 2.3.1 Residual parameter prediction module

Only MRI scans that passed manual QC were used for training of the residual parameter prediction module. For each QC-passed scan, affine parameters were randomly sampled to generate a synthetic misalignment, with a combination of rotation, scaling, and translation sampled from the ranges (−5°, 5°), (0.85, 1.15), and (−7, 7) mm, respectively. The resulting affine transformation was concatenated with the original affine transformation and applied to the MRI scan in native space to generate a synthetically misaligned image in ICBM space. As shown in Supplementary Fig S1, the range of synthetic misalignment covers the full range of misalignment observed in real QC-failed scans. A 3D ResNet18 was implemented and trained on synthetically misaligned 3D MRI volumes to predict the residual affine parameters (Fig. 2A). The 3D architecture captures voxel-level spatial information, while the residual connections facilitate network optimization.

#### 2.3.2 Classification Module

The classification module includes two approaches: 1. RACOON-P: an SVM classifier using the predicted residual affine parameters as features; 2. RACOON-C: a fine-tuned deep learning classifier using the 3D T1w volume as input. Both classifiers predict a binary QC label (pass = 1; fail = 0). The deep learning classifier was initialized using the pretrained ResNet18 backbone from the residual parameter prediction module (Fig. 2B). Additional residual blocks and an MLP classification head were added to the pretrained backbone. The network was first fine-tuned on real QC-passed and QC-failed MRI scans from Dataset C. Details for Dataset C are available in Supplementary Table S3.

#### 2.3.3 Correction Module

The correction module uses the residual affine parameters predicted by the residual parameter prediction module to recover QC-failed MRI scans to a QC-passed alignment (Fig. 2C). Inspired by the iterative registration framework proposed by Ma et al., we also designed a multi-stage inference strategy (Ma et al., 2024). Assuming *T_ori_* denotes the original affine transformation and 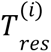 is the predicted residual transformation for the *i*-th inference, the corrected affine transformation for the *i*-th iteration is 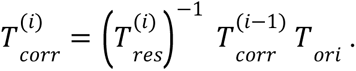 The iterative process is terminated when the predicted residual misalignment distance at two consecutive stages falls below a predefined threshold (δ = 10^−4^ *mm*).

### 2.4 Model Training

#### 2.4.1 Data Processing

To improve model generalizability, no preprocessing was performed for the images that were used to train and validate the performance of RACOON. The linear registration transformations that were used for training were however derived based on PELICAN, a validated longitudinal MRI processing pipeline normalization (Dadar et al., 2025), which performs the following preprocessing steps prior to linear registration. Each scan is denoised and corrected for intensity non-uniformity, and intensity normalized before being registered to the MNI-ICBM152-2009c template (V. Fonov et al., 2011; Manera et al., 2019). The resulting transformations from PELICAN were then applied to raw unpreprocessed images in the native space to derive the set of images that were used to train RACOON. Both synthetic and real MRI scans were center-cropped to a fixed size of 192 × 224 × 192 voxels before network input. Image intensities were normalized to the range [0, 1].

#### 2.4.2 Augmentation

During training, no augmentation was applied to 50% (randomly selected) of the MRI scans. For the remaining 50%, Rician noise, bias-field inhomogeneity, and defacing were randomly applied in both the residual parameter prediction and classification modules. Bias-field augmentation was performed using RandomBiasField from TorchIO, with coefficients sampled from the range [−1, 1] (Pérez-García et al., 2021). Defacing augmentation was included to improve model robustness to different defacing procedures used across datasets, such as UK Biobank (Alfaro-Almagro et al., 2018) and CamCAN. Many defacing methods transform facial masks from template space to the native space for defacing (Bischoff-Grethe et al., 2007a; Cox, 1996; Faruk Gulban et al., n.d.). Following this approach, nine facial masks were generated: N = 4 were manually segmented in the stereotaxic space based on randomly selected cases and N = 5 automated masks were created using five commonly used defacing methods (AFNI, FreeSurfer, FSL, pydeface, and mridefacer)(Bischoff-Grethe et al., 2007b; Buimer et al., 2021; Cox, 1996). During defacing augmentation, one of the nine masks was randomly selected and registered to the native space in an individual MRI scan with SimpleITK. Examples of the facial masks and corresponding defaced MRI scans are shown in Supplementary Fig S2.

#### 2.4.3 Loss function

For the residual parameter prediction module, the target affine parameters were rescaled before model training to balance the contributions of rotation, scaling, and translation to the loss and improve the model’s sensitivity to different types of registration errors. Rotation parameters were left unchanged, whereas the scale and translation parameters were rescaled as follows: 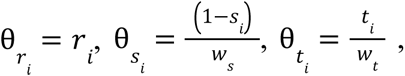 where *r_i_*, *s_i_*, *t_i_* are the ground-truth residual rotation, translation, and scale parameters along spatial axis *i*, respectively. *w_s_* and *w_t_* are weighting factors for scale and translation (see details in section 2.4.4). The 3D ResNet18 was optimized using an L2 loss between the predicted and target affine parameters in the rescaled parameter space, as follows:

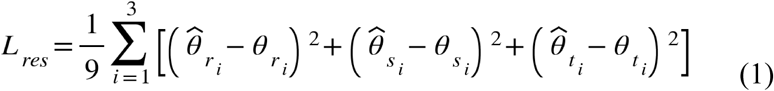

For classification module, the model was trained or fine tuned with binary cross-entropy loss:

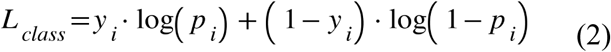

where *y_i_* denotes the ground-truth visual QC label for the i-th scan (fail = 0; pass = 1), *p_i_* denotes the predicted probability of passing QC from RACOON.

#### 2.4.4 Model Configuration and hyperparameter selection for the residual parameter prediction module

Three experiments were conducted to determine the model configuration and training parameters for the residual parameter prediction module. First, a template-input ablation experiment was performed using Dataset A (Supplementary Table S1) to evaluate whether including the template image improved model performance. Three input configurations were compared: no template, a static template, and a moving template where extra transformations were applied to the template and the parameters estimated by the model needed to transform the image from native space to the moved template. The scaling and translation weighting factors, *ws* and *w_t_*, were initially set to 80 and 3.0, respectively. Model performance was evaluated using five-fold cross-validation.

Second, a grid search was performed to determine the optimal weighting factors for scaling and translation with *w_s_* ∈ {50, 60, 70, 80, 90, 100} and *w_t_* ∈ {0.8, 0.9, 1.0, 1.5, 2.0, 2.5, 3.0}. These experiments were performed using Dataset A (Supplementary Table S1) with the model configuration without a template input.

Finally, the effect of training sample size was evaluated using Dataset B (Supplementary Table S2). Five datasets were reserved as independent test sets. For the remaining eight datasets, 20% of the MRI scans were randomly selected for validation, while the remaining scans formed the training pool. Training subsets ranging from 4,000 to 15,000 scans, in increments of 1,000, were sampled from this pool. Five-fold cross-validation was performed for each training sample size.

#### 2.4.5 Implementation and training details

A dropout rate of 0.2 was applied to the first MLP layer. The model was trained with a batch size of 60 using the AdamW optimizer with an initial learning rate of 1.9×10^−5^. A ReduceLROnPlateau scheduler was used to reduce the learning rate by half if the validation loss did not improve for five consecutive epochs, with a scheduler threshold of 0.001. Early stopping was triggered if no improvement was observed for 10 consecutive epochs, and the maximum number of training epochs was set to 90. Automatic mixed-precision training was implemented using PyTorch’s torch.cuda.amp.autocast to reduce memory usage and improve computational efficiency.

### 2.5 Performance Evaluation

For the residual parameter prediction module, the predicted parameters were first transformed back to their original scales using the corresponding weighting factors. The mean absolute error (MAE) between the predicted and ground-truth parameters was calculated separately for rotation, scaling, and translation. To quantify the overall spatial misalignment, we calculated the root squared mean distance (RMSE) consistent with previous studies (de Senneville et al., 2020; V. S. Fonov et al., 2022):

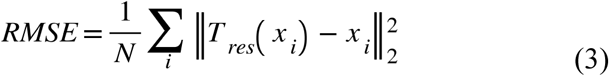

where N is the number of sampled points, *x_i_* denotes the spatial coordinate of the i-th point, and 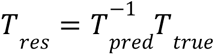 represents the residual transformation between the predicted and ground-truth affine transformations. Six predefined points located near the brain boundary in ICBM template space were used to quantify RMSE with the following MNI coordinates: (0,−105,15), (−70,−21,18), (70,21,18), (0,73,15), (0,-20,83), and (0,-20,-47) mm. These points approximately represent the anterior, left, right, posterior, superior, and inferior extents of the brain, respectively.

For the classification module, 10% of Dataset C was held out as an independent test set, while the remaining 90% was used for five-fold cross-validation. All five trained models, together with the publicly available RegQC and DARQ models, were evaluated on the same held-out test set. Classification performance was assessed using accuracy, sensitivity, specificity, precision, F1 score, and area under the receiver operating characteristic curve (AUC).

### 2.6 Assessing the variability of QC-passed registrations

To estimate the level of variability that can be expected in affine parameters for QC-passed registrations, linear registration was performed and QCed twice on three datasets (ADNI, UKBB, and PPMI). Only MRI scans that passed QC in the two runs were included in this subsequent analysis. For each affine parameter, a linear regression model was fitted using the parameter from the first registration run to predict the parameter from the second run. The MAE between the predicted and observed parameters in the second run was then calculated. In addition, the RMSE between the two affine matrices obtained from the two registration runs was calculated as:

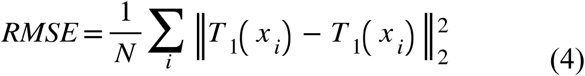

where *T*_1_ and *T_2_* represent the affine transformations obtained from the first and second registration runs, respectively. The same six predefined points in ICBM template space described in Section 2.5 were used to get the RMSE between two linear registration runs. The RMSE between the two QC-passed runs represent the minimum detectable misalignment error we can achieve in both residual parameter prediction module and correction module.

### 2.7 Data and Code Availability

Detailed information for the publicly available datasets used in this study are provided in Supplementary S1. PELICAN image processing pipeline is open source and available at: https://zenodo.org/records/17168419. The source codes, trained models, and instructions for running RACOON are publicly available at https://github.com/ZhaojinChen/RACOON.

## 3. Results

### 3.1 Variability of real QC-passed images

The variability of QC-passed registrations was assessed using the MAE of the affine parameters and the RMSE between repeated registration runs (Table 3). ADNI showed the largest MAE across most affine parameters and the highest RMSE (0.786 mm) among the three datasets, indicating greater variability between repeated registrations. A consistent axis-specific pattern was also observed across all three datasets, with the largest rotation MAE along the x-axis and the lowest scaling and translation MAEs along the x-axis.

**Table 3.** Performance of different template-input configurations on the test sets of Dataset A across five folds. (mean ± standard deviation). The best performance for each affine parameter and RMSE is highlighted in bold.

| Methods | Axis | Rotation(°) | Scale | Translation(mm) | RMSE (mm) |
| --- | --- | --- | --- | --- | --- |
| static template | X | 0.530±0.096 | 0.009±0.001 | 0.859±0.152 | 2.285±0.420 |
|  | Y | 0.454±0.121 | 0.010±0.002 | 0.968±0.100 |  |
|  | Z | 0.424±0.092 | 0.013±0.007 | 0.829±0.140 |  |
| without template | X | **0.484±0.031** | **0.009±0.0007** | **0.782±0.034** | **2.062±0.122** |
|  | Y | **0.421±0.052** | **0.009±0.0005** | **0.915±0.036** |  |
|  | Z | **0.397±0.038** | **0.010±0.0008** | **0.721±0.055** |  |
| Moving template | X | 0.584±0.061 | 0.010±0.001 | 0.863±0.030 | 2.384±0.088 |
|  | Y | 0.460±0.049 | 0.010±0.0006 | 1.051±0.047 |  |
|  | Z | 0.419±0.032 | 0.012±0.0004 | 0.922±0.063 |  |

### 3.2 Residual Parameter Prediction module

#### 3.1.1 Template input ablation and weighting factor selection

Fonov et al. previously showed that including a template image did not improve classification performance when all registered T1w MRI scans were already in the same template space (V. S. Fonov et al., 2022). We further evaluated whether including the template as an additional input could improve residual parameter prediction (Table 3). The model without a template achieved the lowest mean RMSE (2.062 mm) and the lowest MAE across the affine parameters. In comparison, the model using a moving template resulted in the highest mean RMSE (2.384 mm). Based on these results, the model without a template input was selected for the subsequent experiments.

The performance of models trained with different scaling and translation weighting factors was evaluated (Fig. 3). The model with model configuration 70/2.5 (scaling weighting factor: 70; translation weighting factors: 2.5) achieved an RMSE of 1.392 mm on the validation set and the lowest RMSE of 1.435 mm on the independent test sets. This configuration also showed lowest translation MAE, ranging from 0.43 to 0.65 mm in test sets as shown in Supplementary Fig S3. Therefore, the 70/2.5 configuration was selected for the subsequent experiments.

**Fig 3.**
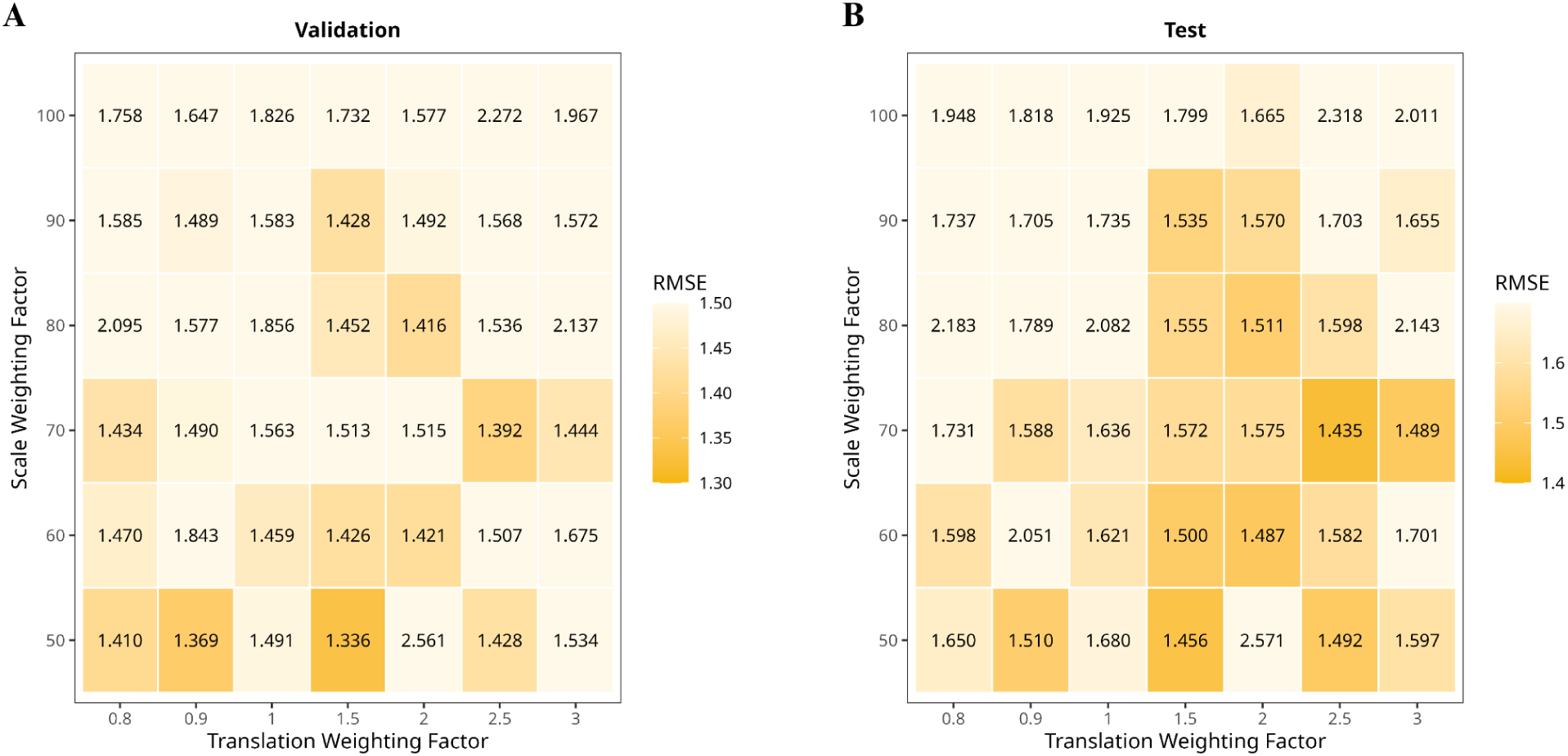
Heatmaps of mean RMSE for different combinations of scaling and translation weighting factors across five folds in Dataset. **A.** (A) Validation set performance. (B) Independent test set performance. Darker colors indicate lower RMSE and better performance.

#### 3.1.2 Training sample size and final model performance

The effect of training sample size was evaluated by training models with sample sizes ranging from 4,000 to 15,000 and assessing their performance on the validation and independent test sets across five folds (Fig. 4). RMSE showed an overall decreasing trend as the training sample size increased from 4,000 to 13,000. The lowest mean RMSE on both the validation and independent test sets was observed at a training sample size of 13,000. Similarly, as shown in Supplementary Fig. S4, the mean MAE across the five folds for the affine parameters was also lowest at a sample size of 13,000 in test sets. Increasing the training sample size to 14,000 or 15,000 resulted in comparable or slightly higher RMSE, suggesting that model performance reached a plateau at approximately 13,000 training samples. Therefore, a training sample size of 13,000 was selected for the final residual parameter prediction model.

**Fig 4.**
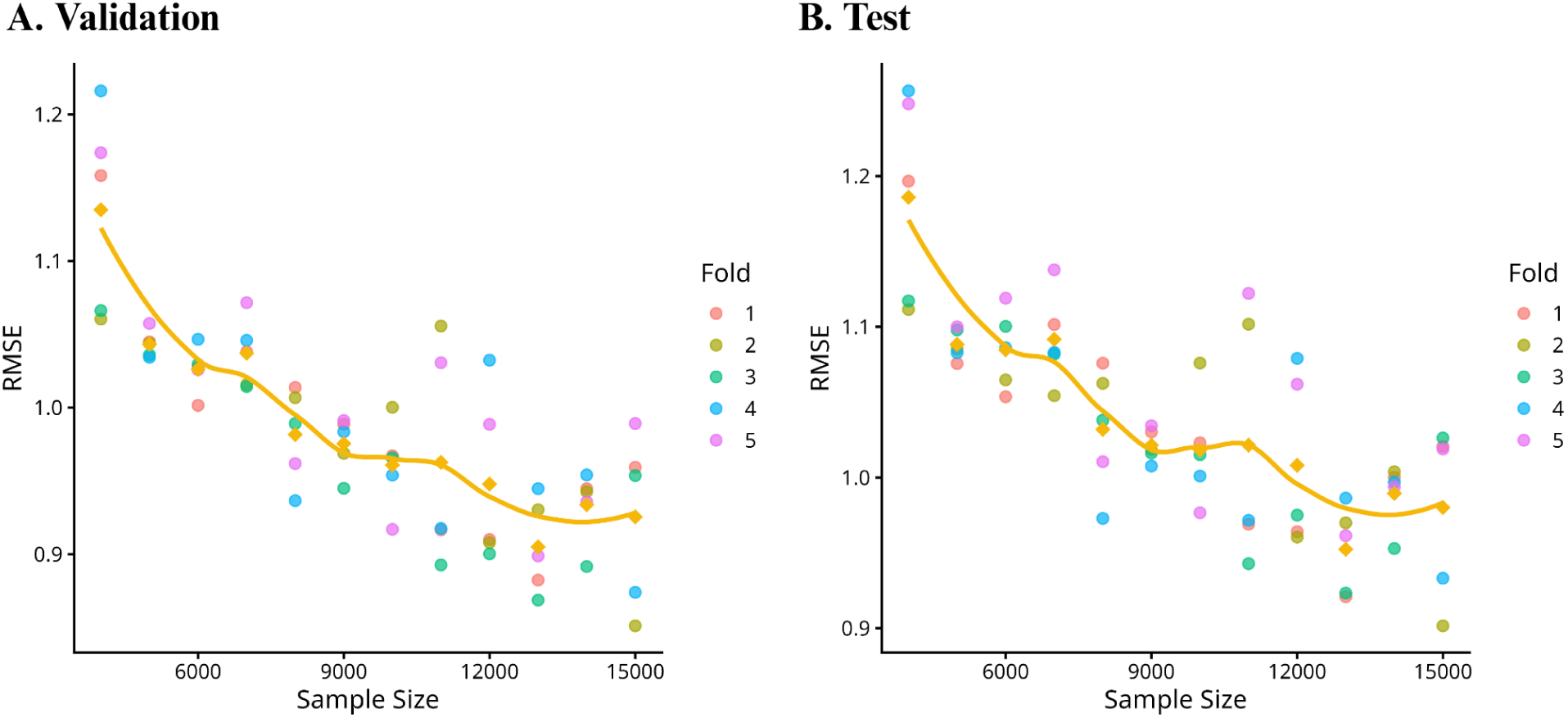
RMSE of the residual parameter prediction models trained with different sample sizes on (A) the validation set and (B) the independent test sets of Dataset. **B.** Colored points represent the mean RMSE across all MRI scans within each of the five folds, and the yellow line represents the mean RMSE across the five folds at each training sample size.

During the inference stage, the predicted affine parameters were first averaged across the five folds, and the RMSE was then calculated using the averaged predicted parameters. For the model trained with 13,000 samples, the RMSE on test sets of Dataset B was 0.861 ± 0.299 mm, indicating submillimeter residual misalignment. Performance across the five independent test datasets is shown in Fig. 5. The predicted parameters were closely distributed around the identity line across all nine affine parameters, with comparable performance across datasets. Among the five datasets, NIFD generally showed the largest MAE for all the affine parameters, but still achieved a submillimeter RMSE of 0.92 ± 0.35 mm, comparable to the overall performance. An axis-specific pattern was also observed across datasets, with higher rotation MAE along the x-axis but lower scaling and translation MAE along the x-axis compared with the other two axes. Specifically, the x-axis rotation MAE ranged from 0.305° to 0.336° across datasets, whereas the y- and z-axis rotation MAEs were approximately 0.19°–0.23°. In contrast, the x-axis translation MAE ranged from 0.091 to 0.139 mm, compared with approximately 0.20–0.26 mm along the y- and z-axes.

**Fig 5.**
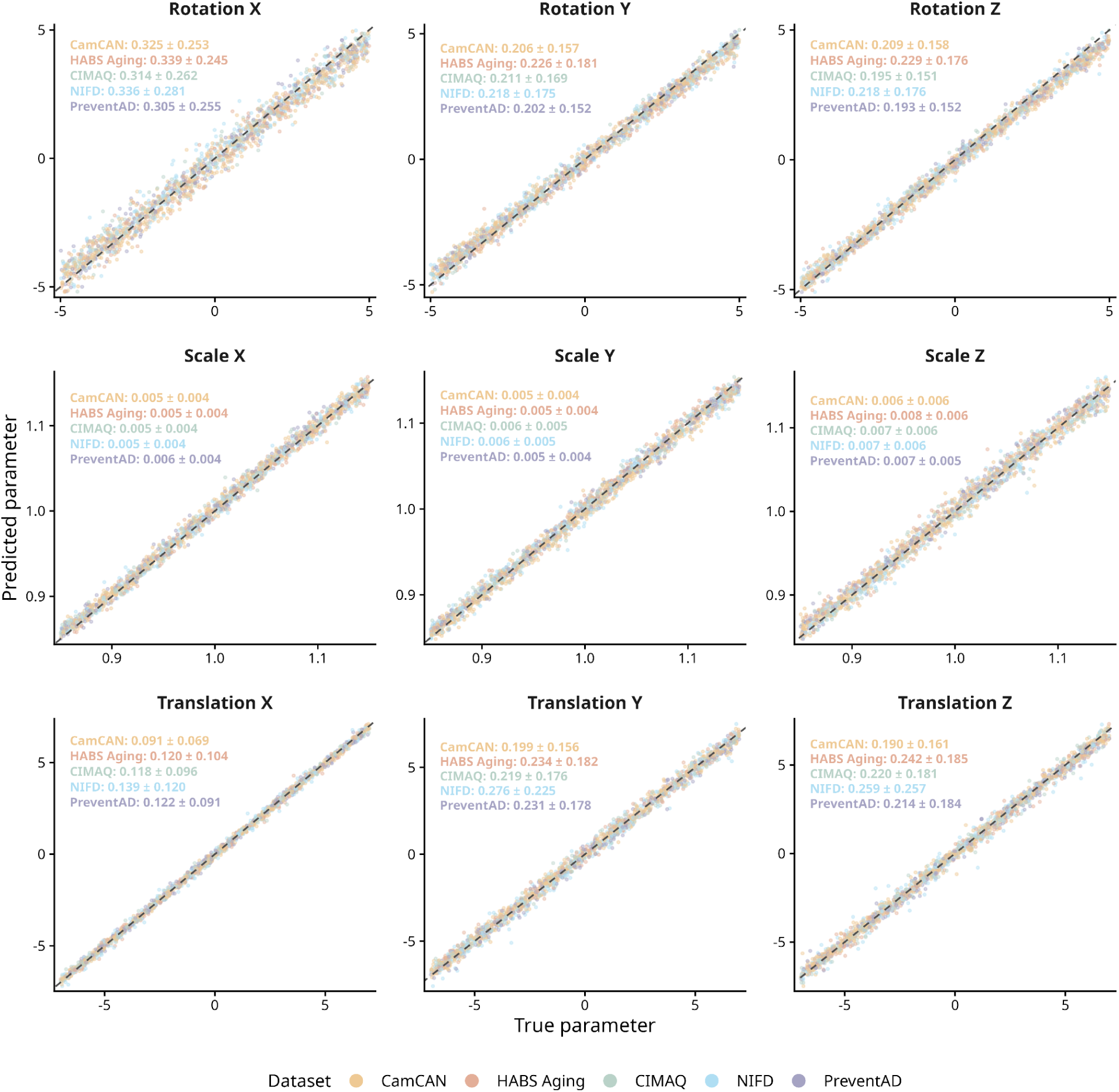
Predicted versus ground-truth affine parameters on the five independent test datasets for the model trained with 13,000 samples from Dataset. **B.** Each point represents an MRI scan and is colored according to the dataset. The black dashed line represents the identity line, where the predicted parameter equals the ground-truth parameter. Values in each panel indicate the MAE (mean ± standard deviation) for each dataset.

Iterative inference further improved model performance, achieving an RMSE of 0.778 ± 0.286 mm (t=-8.36, p<0.001). As shown in Fig. 6, the MAE of all affine parameters decreased across the five independent test datasets. The improvement was particularly evident for rotation, with MAE decreasing by approximately 0.1°–0.4° compared with single-stage inference.

**Fig 6.**
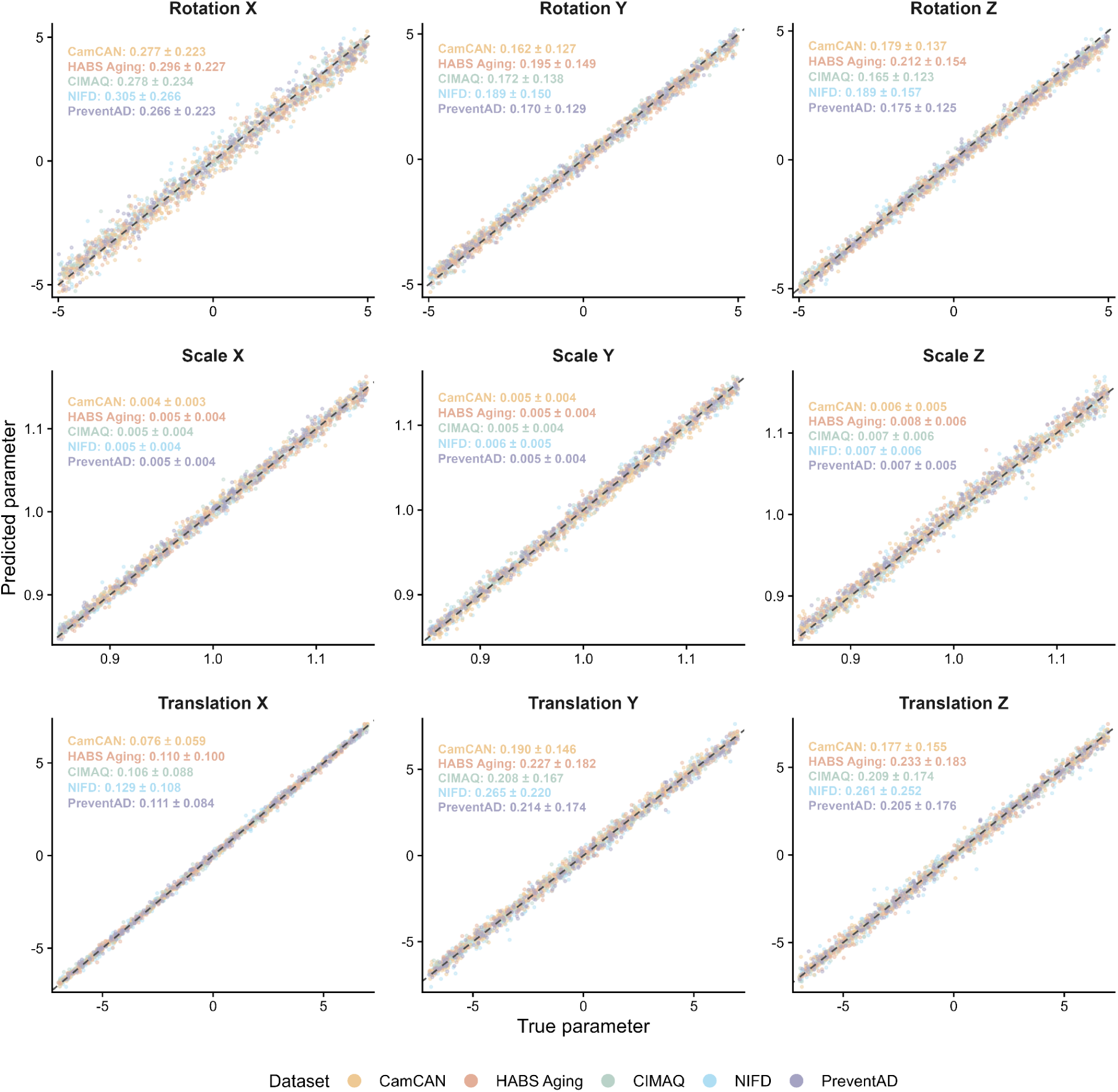
Predicted versus ground-truth affine parameters on the five independent test datasets using iterative inference.

### 3.3 Classification Module

To obtain comparable results, RegQCNET was evaluated on the same test set, and the optimal RMSE threshold was determined using ROC curve analysis. The optimal threshold for RegQCNET was 5.48 mm, with RMSE values of 3.88 ± 1.63 mm and 4.31 ± 2.00 mm for QC-passed and QC-failed scans, respectively. As shown in Table 4, RACOON-C achieved the highest classification accuracy (0.767), while RACOON-P achieved the second-highest accuracy (0.721), followed by direct classification, DARQ, and RegQCNET. RACOON also showed higher precision than the other methods, with RACOON-C achieving the highest precision (0.744).

**Table 4.** Classification performance of different methods on the held-out test set of Dataset D, averaged across five folds. RACOON-P uses the residual affine parameters predicted by the residual parameter prediction module as input to an SVM with an RBF kernel. RACOON-C is a 3D ResNet18 classifier fine-tuned from the residual parameter prediction module.

| Method | Accuracy | Balanced accuracy | Precision | Recall | AUC | Confusion matrix [tn, fp: fn, tp] |
| --- | --- | --- | --- | --- | --- | --- |
| DARQC | 0.545 | 0.551 | 0.521 | 0.799 | - | [197, 454; 124, 494] |
| RegQCNet | 0.523 | 0.564 | 0.475 | 0.886 | - | [101, 317; 37, 287] |
| Resnet 18 | 0.719±0.006 | 0.721±0.005 | 0.686±0.034 | 0.798±0.083 | 0.804±0.009 | [479.2, 264.8; 144.4, 568.6] |
| RACOON-P | 0.721±0.002 | 0.723±0.002 | 0.673±0.003 | 0.838±0.003 | 0.801±0.002 | [453.4, 290.6; 115.80, 597.20] |
| RACOON-C | **0.767±0.006** | **0.768±0.005** | **0.744±0.02** | **0.803±0.003** | **0.853±0.008** | **[545.4, 198.6; 140.6, 572.6]** |

### 3.4 Correction Module

The MRI scans identified as QC failures by the classification module were corrected using the predicted residual affine parameters. Fig. 7A shows an example, where the QC-failed MRI scan registered by ANTs exhibited a scaling error across all three views and a rotation error that was most apparent in the axial view. RACOON successfully corrected these misalignments, resulting in improved agreement between the MRI scan and the template outlines across all views (Fig. 7B). The corrected scan subsequently passed visual QC (blind to the correction status).

**Fig. 7.**
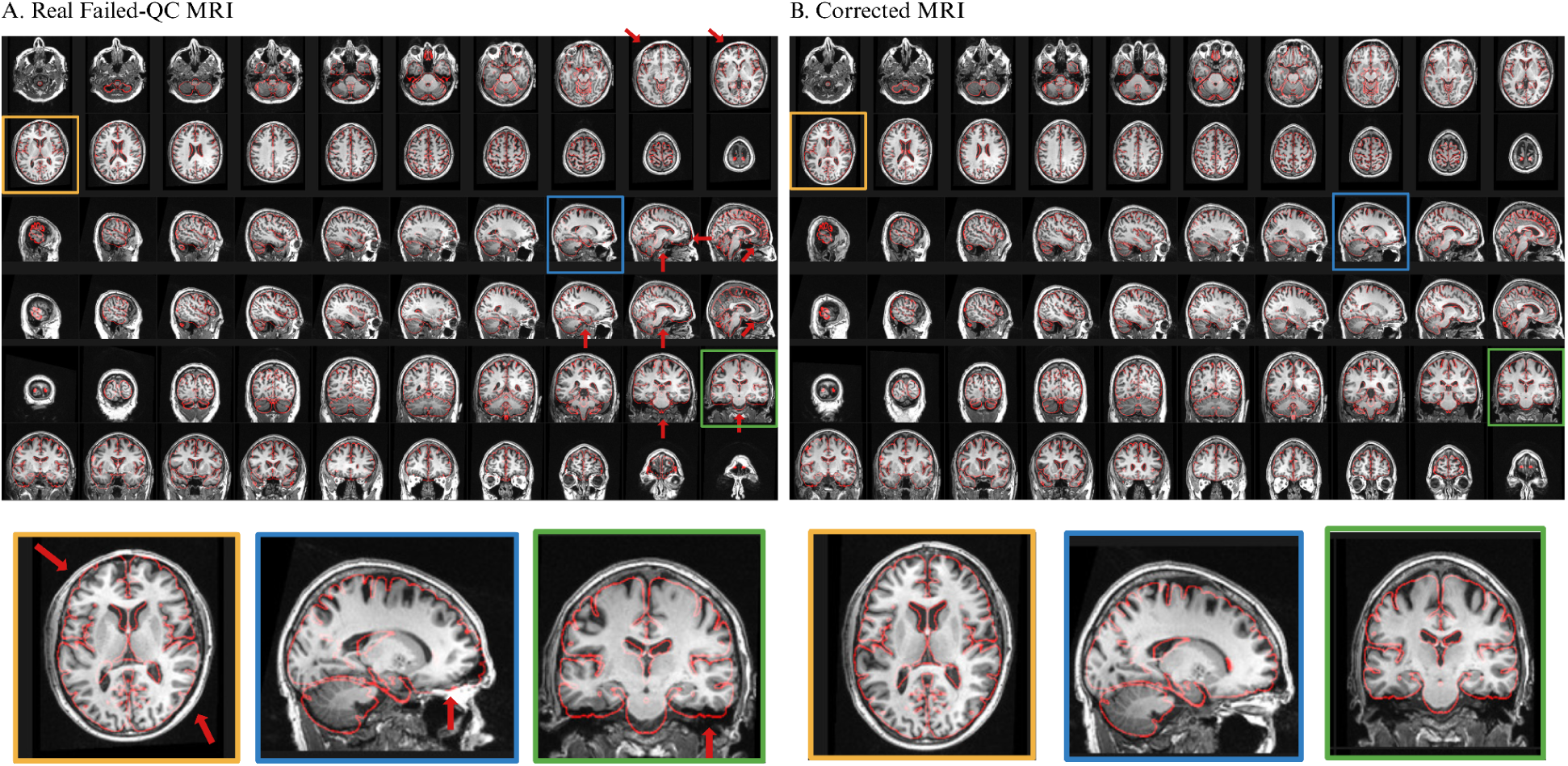
Examples of MRI scans that failed QC after ANTs registration and were successfully corrected by RACOON. The red contours represent the template outline, and the red arrows in A indicate regions of registration misalignment. Selected slices are enlarged and shown below the corresponding images for better visualization.

## 4. Discussion

In this work, we proposed RACOON, a framework for accurate automatic quality control and misalignment correction of stereotaxic linear registration. We demonstrated that RACOON can detect submillimeter misalignment, identify QC-failed MRI scans with subtle residual misalignment, and correct real registration failures.

Most previous deep learning-based registration methods use image similarity metrics, such as mutual information (MI) or normalized cross-correlation (NCC), as loss functions to maximize the intensity similarity between the template and individual MRI scans (De Vos et al., 2019; Dubost et al., 2020; Hoffmann et al., 2021; Iglesias, 2023). However, similarity-based registration has several limitations. First, MRI scans with substantial pathological changes are generally more challenging to register, particularly when the reference template is constructed from a relatively healthy population. Second, previous deep learning-based registration methods used the same or similar image similarity loss functions as conventional optimization-based methods. Therefore, although they can substantially reduce registration time, they may converge to similar registration solutions and have generally shown comparable or suboptimal performance compared with conventional optimization-based methods such as ANTs (Avants et al., 2009). More importantly, maximizing image similarity does not necessarily allow the model to learn the range of alignments considered acceptable by human raters. Based on these considerations, RACOON was designed to directly predict residual affine parameters rather than optimize an image similarity-based loss.

Several model configurations, including the scaling and translation weighting factors, template input, and training sample size, were systematically evaluated to determine the final model configuration for the residual parameter prediction module. With iterative inference, the final model achieved an RMSE comparable to the variability observed between repeated QC-passed registrations in ADNI. This suggests that the residual misalignment after correction is approaching the level of variability observed among registrations that are considered acceptable by visual QC. Similar to Ma’s study (Ma et al., 2024), iterative inference reduced the RMSE from 0.861 mm to 0.778 mm at the cost of increased inference time. As shown in Fig. S5, prediction errors were larger for scans with greater initial misalignment, with a strong correlation observed between the prediction error and the ground-truth parameters (r = -0.53 for rotation along z axis). Iterative inference reduced this dependence, resulting in reduced correlation values between prediction errors and ground-truth values across all affine parameters (r = -0.37 for rotation along z axis).

Previous studies have shown that defacing can introduce subtle variations in estimated affine transformation parameters (Schwarz et al., 2021). RegQCNET reported that intensity non-uniformity can slightly reduce the accuracy of registration error prediction(de Senneville et al., 2020). Based on these observations, both defacing and bias-field augmentation were included during model training to improve robustness to variations in image preprocessing and intensity distributions. Incorporating defacing augmentation reduced this generalization bias when the model was applied to an independent defaced dataset. The model trained without defacing augmentation achieved an RMSE of 1.39 ± 0.60 mm on the validation set, which increased significantly to 1.69 ± 0.56 mm on the defaced CamCAN dataset (t=-11.68, p<0.001). In contrast, after defacing augmentation, no significant difference in RMSE was observed between the validation and CamCAN datasets, with RMSEs of 1.04 ± 0.37 mm and 1.03 ± 0.31 mm, respectively.

Interestingly, a consistent axis-specific pattern was observed in both repeated QC-passed registrations and the synthetic test data, with lower variability and prediction error for scaling and translation along the x-axis, but higher variability and prediction error for rotation. This pattern may reflect differences in anatomical variability, image resolution, and head positioning across spatial directions. Similar to our findings, Zhang et al. also reported direction-dependent differences in registration accuracy for arterial spin labeling MRI, with larger rotation prediction errors along the x-axis, suggesting that registration uncertainty may not be uniform across spatial directions (Zhang et al., 2023). In particular, rotation around the x-axis primarily affects the orientation of the brain in the y–z plane and may be more sensitive to variations in head positioning, such as differences in neck flexion and head support during MRI acquisition. Anatomical changes, such as asymmetric ventricular enlargement, may further increase the difficulty of achieving consistent alignment in the y–z plane and contribute to greater variability in rotation around the x-axis (Fig S6). In addition, as shown in Table 1, some datasets had lower spatial resolution along the x-axis than along the other two axes. This anisotropic resolution may contribute to the higher variability observed for scaling and translation along the x-axis as well as the lower bound accuracy achievable based on registration and re-registration RMSE error. Overall, the observed axis-specific pattern is likely influenced by multiple factors, including image acquisition, head positioning, and alignment of certain anatomical structures. The presence of a similar axis-specific pattern in both real registration variability and synthetic prediction errors suggests that the observed prediction error may partly reflect the intrinsic variability of affine registration rather than a model-specific bias.

For the classification module, RACOON-C achieved a balanced accuracy of approximately 76.8%, representing an improvement of approximately 4.8 percentage points over direct classification, despite the subtle differences between QC-passed and QC-failed scans. In addition, RACOON-C improved precision by 6.8 percentage points compared with direct classification and correctly identified more true registration failures. More accurate identification of registration failures can reduce unnecessary data exclusion while preventing misregistered scans from entering downstream analyses, thereby improving data retention and the reliability of subsequent analyses. RACOON-C uses the pretrained backbone from the residual parameter prediction module rather than training the network from scratch. This allows the classification model to use both anatomical features from the 3D MRI volume and features that have already been optimized to characterize residual registration errors, which may contribute to its higher accuracy compared with both RACOON-P and direct classification. As shown in Fig. S7, the RMSE distributions estimated by both RegQCNET and RACOON showed substantial overlap between QC-passed and QC-failed scans. Importantly, the mean RMSE of the real QC-failed scans was lower than the thresholds used for RegQCNET (10 mm) and DARQ (de Senneville et al., 2020; V. S. Fonov et al., 2022), suggesting that many failures in our dataset represented relatively subtle registration errors. This may partly explain the lower performance of RegQCNET and DARQ on our dataset. Nevertheless, RACOON-P achieved performance comparable to direct classification using only the predicted residual affine parameters. This finding suggests that the predicted parameters themselves contain meaningful information about registration quality and could potentially be used by other groups to define their own QC criteria according to their datasets and processing pipelines.

An important strength of RACOON is that the residual parameter prediction module was trained using 13,000 manually QC-passed MRI scans from 11 datasets, allowing the model to learn population-level patterns to achieve QC-passed alignments for misaligned scans. Manual QC was performed by an experienced rater who has evaluated more than 100,000 cases, providing a highly consistent set of QC-passed registrations with low variability in affine parameters. Following systematic optimization of the model configuration, RACOON achieved a residual misalignment approaching the level of variability observed among registrations considered acceptable by visual QC.

Most importantly, unlike methods that focus only on identifying registration failures, RACOON provides modules for both the identification and correction of registration errors. The classification module first identifies QC-failed scans, which can then be processed by the correction module to reduce their residual misalignment. This may allow scans that would otherwise be excluded because of registration failure to be retained for downstream analyses, thereby reducing data loss and preserving study sample representation.

Several limitations warrant mentioning. Compared with DARQ, the classification module of RACOON was mainly trained using linear registrations generated by ANTs. Based on our experience and according to our previous study (Dadar et al., 2025), ANTs shows robust performance for linear registration even in scans with poor initial head positioning. Thus, PELICAN pipeline was developed based on registrations generated by ANTs. As a result, the types and ranges of registration errors in the real QC-failed scans may be limited, however, we believe that our initial residual estimation module explores the parameter space rather exhaustively and is tool-agnostic and addresses this issue. Nevertheless, future work can include failed registrations generated by other linear registration methods to improve the robustness of RACOON across different registration methods. In addition, the transformation parameters used to train RACOON did not include shearing. Therefore, registration failures caused by shearing are not corrected by the current correction module (see Fig. S8 for an example). Future work could incorporate shearing parameters to address this limitation. RACOON currently focuses only on the QC and correction of linear registration. Future work could extend RACOON into an integrated quality control framework that evaluates multiple preprocessing steps, such as nonlinear registration, skull stripping, and tissue segmentation. Similar to DARQ (V. S. Fonov et al., 2022), RACOON was developed and validated using the MNI-ICBM152-2009c template. As such, retraining would be required for use in applications where the target template substantially differs from MNI-ICBM152-2009c stereotaxic alignment. However, given that most other average templates (Dadar et al., 2021, 2022; V. Fonov et al., 2011; V. S. Fonov et al., 2009) have also been defined in the same stereotaxic space and have similar brain boundaries, RACOON would also be applicable for evaluating linear registrations to other templates in the same stereotaxic space.

Overall, we demonstrated that RACOON can correct linear registration errors and accurately identify subtle registration errors compared to state-of-arts methods. The residual parameter prediction module showed consistent performance across datasets, scanners, and different defacing techniques, demonstrating its robustness to variations in image acquisition and preprocessing. Finally, the error correction module available in RACOON is an invaluable addition which can further reduce biases introduced by failed linear registration in aging and neurodegenerative disorders cohort studies. RACOON therefore provides an automated, fast and robust solution for both quality control and correction of linear registration in large-scale neuroimaging studies.

## 5. Author Contribution

Z.C.: Coding, study design, data analysis, and manuscript writing; R.M.: Quality control; A.M.: Quality control; W.A.-G.: Study design and project supervision; Y.Z.: Study design, conceptualization, manuscript writing, project supervision, funding; M.D.: Study design, conceptualization, manuscript writing, project supervision, funding.

## 6. Acknowledgments

The authors gratefully acknowledge all the open datasets that made this work possible: UK biobank (Sudlow et al., 2015), The Alzheimer’s Disease Neuroimaging Initiative (Weiner et al., 2010), Parkinson’s Progression Markers Initiative (Marek et al., 2011), Harvard Aging Brain(Dagley et al., 2017), The National Alzheimer’s Coordinating Center(Beekly et al., 2004), FTLDNI database and grant (R01 AG032306), Canadian Consortium on Neurodegeneration in Aging(Chertkow et al., 2019), The Consortium for the early identification of Alzheimer’s disease–Quebec(Belleville et al., 2019), ALLFTD Consortium(Heuer et al., 2024), PreventAD(Villeneuve et al., 2025), Human Brain Project(Bookheimer et al., 2019; Van Essen et al., 2013), Cambridge Centre for Ageing and Neuroscience (Cam-CAN) study(Shafto et al., 2014). The authors also acknowledge Digital Research Alliance of Canada (https://www.alliancecan.ca/en) for the usage of the computing resources in the current work.

## 9. Ethics

Ethics approval for all datasets used was obtained at respective sites.

## Reference

Alfaro-Almagro, F., Jenkinson, M., Bangerter, N. K., Andersson, J. L., Griffanti, L., Douaud, G., Sotiropoulos, S. N., Jbabdi, S., Hernandez-Fernandez, M., Vallee, E., & others. (2018). Image processing and Quality Control for the first 10,000 brain imaging datasets from UK Biobank. Neuroimage, 166, 400–424.

Avants, B. B., Tustison, N., Song, G., & others. (2009). Advanced normalization tools (ANTS). Insight j, 2(365), 1–35.

Baron, J.-C., Chételat, G., Desgranges, B., Perchey, G., Landeau, B., de La Sayette, V., & Eustache, F. (2001). In vivo mapping of gray matter loss with voxel-based morphometry in mild Alzheimer’s disease. Neuroimage, 14(2), 298–309.

Beekly, D. L., Ramos, E. M., Van Belle, G., Deitrich, W., Clark, A. D., Jacka, M. E., & Kukull, W. A. (2004). The national Alzheimer’s coordinating center (NACC) database: An Alzheimer disease database. Alzheimer Disease & Associated Disorders, 18(4), 270–277.

Belleville, S., LeBlanc, A. C., Kergoat, M.-J., Calon, F., Gaudreau, P., Hébert, S. S., Hudon, C., Leclerc, N., Mechawar, N., Duchesne, S., & others. (2019). The Consortium for the early identification of Alzheimer’s disease–Quebec (CIMA-Q). *Alzheimer’s & Dementia: Diagnosis*, Assessment & Disease Monitoring, 11(1), 787–796.

Benhajali, Y., Badhwar, A., Spiers, H., Urchs, S., Armoza, J., Ong, T., Pérusse, D., & Bellec, P. (2020). A standardized protocol for efficient and reliable quality control of brain registration in functional MRI studies. Frontiers in Neuroinformatics, 14, 7.

Bischoff-Grethe, A., Ozyurt, I. B., Busa, E., Quinn, B. T., Fennema-Notestine, C., Clark, C. P., Morris, S., Bondi, M. W., Jernigan, T. L., Dale, A. M., & others. (2007a). A technique for the deidentification of structural brain MR images. Human Brain Mapping, 28(9), 892–903.

Bischoff-Grethe, A., Ozyurt, I. B., Busa, E., Quinn, B. T., Fennema-Notestine, C., Clark, C. P., Morris, S., Bondi, M. W., Jernigan, T. L., Dale, A. M., & others. (2007b). A technique for the deidentification of structural brain MR images. Human Brain Mapping, 28(9), 892–903.

Bookheimer, S. Y., Salat, D. H., Terpstra, M., Ances, B. M., Barch, D. M., Buckner, R. L., Burgess, G. C., Curtiss, S. W., Diaz-Santos, M., Elam, J. S., & others. (2019). The lifespan human connectome project in aging: An overview. Neuroimage, 185, 335–348.

Buckner, R. L., Head, D., Parker, J., Fotenos, A. F., Marcus, D., Morris, J. C., & Snyder, A. Z. (2004). A unified approach for morphometric and functional data analysis in young, old, and demented adults using automated atlas-based head size normalization: Reliability and validation against manual measurement of total intracranial volume. Neuroimage, 23(2), 724–738.

Buimer, E. E., Schnack, H. G., Caspi, Y., van Haren, N. E., Milchenko, M., Pas, P., Initiative, A. D. N., Hulshoff Pol, H. E., & Brouwer, R. M. (2021). De-identification procedures for magnetic resonance images and the impact on structural brain measures at different ages. Human Brain Mapping, 42(11), 3643–3655.

Ceccarelli, A., Jackson, J., Tauhid, S., Arora, A., Gorky, J., Dell’Oglio, E., Bakshi, A., Chitnis, T., Khoury, S. J., Weiner, H. L., & others. (2012). The impact of lesion in-painting and registration methods on voxel-based morphometry in detecting regional cerebral gray matter atrophy in multiple sclerosis. American Journal of Neuroradiology, 33(8), 1579–1585.

Chertkow, H., Borrie, M., Whitehead, V., Black, S. E., Feldman, H. H., Gauthier, S., Hogan, D. B., Masellis, M., McGilton, K., Rockwood, K., & others. (2019). The comprehensive assessment of neurodegeneration and dementia: Canadian cohort study. Canadian Journal of Neurological Sciences, 46(5), 499–511.

Cox, R. W. (1996). AFNI: software for analysis and visualization of functional magnetic resonance neuroimages. Computers and Biomedical Research, 29(3), 162–173.

Dadar, M., Camicioli, R., & Duchesne, S. (2022). Multi sequence average templates for aging and neurodegenerative disease populations. Scientific Data, 9(1), 238.

Dadar, M., Fonov, V. S., Collins, D. L., Initiative, A. D. N., & others. (2018). A comparison of publicly available linear MRI stereotaxic registration techniques. Neuroimage, 174, 191–200.

Dadar, M., Manera, A. L., Fonov, V. S., Ducharme, S., & Collins, D. L. (2021). MNI-FTD templates, unbiased average templates of frontotemporal dementia variants. Scientific Data, 8(1), 222.

Dadar, M., Moqadam, R., Metz, A., Chadwick, K., Brzezinski-Rittner, A., & Zeighami, Y. (2025). PELICAN: a longitudinal image processing pipeline for analyzing structural magnetic resonance images in aging and neurodegenerative disease populations. bioRxiv, 2025–09.

Dagley, A., LaPoint, M., Huijbers, W., Hedden, T., McLaren, D. G., Chatwal, J. P., Papp, K. V., Amariglio, R. E., Blacker, D., Rentz, D. M., & others. (2017). Harvard aging brain study: Dataset and accessibility. Neuroimage, 144, 255–258.

de Senneville, B. D., Manjon, J. V., & Coupé, P. (2020). RegQCNET: Deep quality control for image-to-template brain MRI affine registration. Physics in Medicine & Biology, 65(22), 225022.

De Vos, B. D., Berendsen, F. F., Viergever, M. A., Sokooti, H., Staring, M., & Išgum, I. (2019). A deep learning framework for unsupervised affine and deformable image registration. Medical Image Analysis, 52, 128–143.

Dubost, F., de Bruijne, M., Nardin, M., Dalca, A. V., Donahue, K. L., Giese, A.-K., Etherton, M. R., Wu, O., de Groot, M., Niessen, W., & others. (2020). Multi-atlas image registration of clinical data with automated quality assessment using ventricle segmentation. Medical Image Analysis, 63, 101698.

Faruk Gulban, O., Nielson, D., Poldrack, R., Lee, J., Gorgolewski, C., Ghosh, S., & others. (n.d.). Poldracklab/pydeface: V2. 0.0. Zenodo.

Fernandez-Lozano, S., Dadar, M., Morrison, C., Manera, A., Andrews, D., Rajabli, R., Madge, V., St-Onge, E., Shafiee, N., Livadas, A., & others. (2024). QRATER: a collaborative and centralized imaging quality control web-based application. Aperture Neuro, 4, 10–52294.

Fonov, V., Evans, A. C., Botteron, K., Almli, C. R., McKinstry, R. C., Collins, D. L., Group, B. D. C., & others. (2011). Unbiased average age-appropriate atlases for pediatric studies. Neuroimage, 54(1), 313–327.

Fonov, V. S., Dadar, M., Adni, T. P.-A. R. G., & Collins, D. L. (2022). DARQ: Deep learning of quality control for stereotaxic registration of human brain MRI to the T1w MNI-ICBM 152 template. NeuroImage, 257, 119266.

Fonov, V. S., Evans, A. C., McKinstry, R. C., Almli, C. R., & Collins, D. L. (2009). Unbiased nonlinear average age-appropriate brain templates from birth to adulthood. NeuroImage, 47, S102.

Heuer, H. W., Forsberg, L. K., Mester, C. T., Kolander, T., Johnson, N., Brushaber, D., Rosen, H. J., Boeve, B. F., Boxer, A. L., & Consortium, A. (2024). ALLFTD: characterization of Frontotemporal Lobar Degeneration (FTLD) disease trajectories through longitudinal assessment. Alzheimer’s & Dementia, 20, e093231.

Hoffmann, M., Billot, B., Greve, D. N., Iglesias, J. E., Fischl, B., & Dalca, A. V. (2021). SynthMorph: Learning contrast-invariant registration without acquired images. IEEE Transactions on Medical Imaging, 41(3), 543–558.

Iglesias, J. E. (2023). A ready-to-use machine learning tool for symmetric multi-modality registration of brain MRI. Scientific Reports, 13(1), 6657.

Im, K., Lee, J.-M., Lyttelton, O., Kim, S. H., Evans, A. C., & Kim, S. I. (2008). Brain size and cortical structure in the adult human brain. Cerebral Cortex, 18(9), 2181–2191.

Kim, W. H., Ravi, S. N., Johnson, S. C., Okonkwo, O. C., & Singh, V. (2015). On statistical analysis of neuroimages with imperfect registration. Proceedings of the IEEE International Conference on Computer Vision, 666–674.

Ma, T., Zhang, S., Li, J., & Wen, Y. (2024). Iirp-net: Iterative inference residual pyramid network for enhanced image registration. Proceedings of the IEEE/CVF Conference on Computer Vision and Pattern Recognition, 11546–11555.

Manera, A. L., Dadar, M., Collins, D. L., Ducharme, S., Initiative, F. L. D. N., & others. (2019). Deformation based morphometry study of longitudinal MRI changes in behavioral variant frontotemporal dementia. NeuroImage: Clinical, 24, 102079.

Marek, K., Jennings, D., Lasch, S., Siderowf, A., Tanner, C., Simuni, T., Coffey, C., Kieburtz, K., Flagg, E., Chowdhury, S., & others. (2011). The Parkinson progression marker initiative (PPMI). Progress in Neurobiology, 95(4), 629–635.

Markiewicz, C. J., Gorgolewski, K. J., Feingold, F., Blair, R., Halchenko, Y. O., Miller, E., Hardcastle, N., Wexler, J., Esteban, O., Goncavles, M., & others. (2021). The OpenNeuro resource for sharing of neuroscience data. Elife, 10, e71774.

Metz, A., Moqadam, R., Zeighami, Y., Collins, D. L., Villeneuve, S., & Dadar, M. (2025). Quantifying brain atrophy in Frontotemporal Dementia: A head-to-head comparison of neuroimaging techniques. medRxiv, 2025–10.

Miller, K. L., Alfaro-Almagro, F., Bangerter, N. K., Thomas, D. L., Yacoub, E., Xu, J., Bartsch, A. J., Jbabdi, S., Sotiropoulos, S. N., Andersson, J. L., & others. (2016). Multimodal population brain imaging in the UK Biobank prospective epidemiological study. Nature Neuroscience, 19(11), 1523–1536.

Mirhakimi, N., Chatelain, Y., Poline, J.-B., & Glatard, T. (2025). Numerical uncertainty in linear registration: An experimental study. International Workshop on Uncertainty for Safe Utilization of Machine Learning in Medical Imaging, 56–66.

Pérez-García, F., Sparks, R., & Ourselin, S. (2021). TorchIO: a Python library for efficient loading, preprocessing, augmentation and patch-based sampling of medical images in deep learning. Computer Methods and Programs in Biomedicine, 208, 106236.

Raamana, P. R., Theyers, A., Selliah, T., Bhati, P., Arnott, S. R., Hassel, S., Nanayakkara, N. D., Scott, C. J., Harris, J., Zamyadi, M., & others. (2020). Visual QC Protocol for FreeSurfer cortical parcellations from anatomical MRI. BioRxiv, 2020–09.

Shafto, M. A., Tyler, L. K., Dixon, M., Taylor, J. R., Rowe, J. B., Cusack, R., Calder, A. J., Marslen-Wilson, W. D., Duncan, J., Dalgleish, T., & others. (2014). The Cambridge Centre for Ageing and Neuroscience (Cam-CAN) study protocol: A cross-sectional, lifespan, multidisciplinary examination of healthy cognitive ageing. BMC Neurology, 14(1), 204.

Sudlow, C., Gallacher, J., Allen, N., Beral, V., Burton, P., Danesh, J., Downey, P., Elliott, P., Green, J., Landray, M., & others. (2015). UK biobank: An open access resource for identifying the causes of a wide range of complex diseases of middle and old age. PLoS Medicine, 12(3), e1001779.

Thadikemalla, V. S. G., Focke, N. K., & Tummala, S. (2024). A 3d sparse autoencoder for fully automated quality control of affine registrations in big data brain mri studies. Journal of Imaging Informatics in Medicine, 37(1), 412–427.

Van Essen, D. C., Smith, S. M., Barch, D. M., Behrens, T. E., Yacoub, E., Ugurbil, K., Consortium, W.-M. H., & others. (2013). The WU-Minn human connectome project: An overview. Neuroimage, 80, 62–79.

Villeneuve, S., Poirier, J., Breitner, J. C., Tremblay-Mercier, J., Remz, J., Raoult, J.-M., Yakoub, Y., Gallego-Rudolf, J., Qiu, T., Fajardo Valdez, A., & others. (2025). The PREVENT-AD cohort: Accelerating Alzheimer’s disease research and treatment in Canada and beyond. Alzheimer’s & Dementia, 21(10), e70653.

Weiner, M. W., Aisen, P. S., Jack Jr, C. R., Jagust, W. J., Trojanowski, J. Q., Shaw, L., Saykin, A. J., Morris, J. C., Cairns, N., Beckett, L. A., & others. (2010). The Alzheimer’s disease neuroimaging initiative: Progress report and future plans. Alzheimer’s & Dementia, 6(3), 202–211.

Zhang, Z., Yang, H., Guo, Y., Bolo, N. R., Keshavan, M., DeRosa, E., Anderson, A. K., Alsop, D. C., Yin, L., & Dai, W. (2023). Affine image registration of arterial spin labeling MRI using deep learning networks. NeuroImage, 279, 120303.

