## Supplemental Materials for "Automatic Quality Control and Error Correction in MRI linear registration via a Residual Parameter Prediction Network for T1w MRI"

**S1. detailed information for datasets used in this study**

UK Biobank: It is a large population-based cohort that recruited participants aged 40–69 years. Compared with disease-specific clinical cohorts, UK Biobank represents a relatively healthy population. The released T1w MRI scans were defaced using FSL. To balance the effects of defacing and sample size across datasets, we included 15,742 T1w MRI scans from UK Biobank in this study.

ADNI: The Alzheimer's Disease Neuroimaging Initiative aims to develop biomarkers for the detection and longitudinal tracking of Alzheimer's disease progression. Compared with other cohorts, ADNI is a large multicenter study with MRI data acquired across multiple sites, scanners, and magnetic field strengths. In this study, we primarily included 388 T1-weighted MRI scans from the ADNI1 cohort.

PPMI: The Parkinson's Progression Markers Initiative is a longitudinal, multicenter study designed to identify and validate biomarkers associated with the onset and progression of Parkinson's disease (PD). Although PPMI provides longitudinal follow-up data, only cross-sectional T1-weighted MRI scans were included in this study. In total, 3,133 MRI scans were included.

HABS-Aging: The Harvard Aging Brain Study focuses on understanding the transition from cognitively normal aging to Alzheimer’s disease (AD), with particular emphasis on how amyloid and tau pathology are associated with subsequent cognitive decline. Unlike ADNI, most HABS participants were cognitively normal at baseline, enabling the investigation of brain changes during the preclinical stages of AD. To avoid overrepresenting participants with multiple longitudinal scans, we randomly selected one MRI scan per participant from the longitudinal cohort. A total of 278 T1-weighted (T1w) MRI scans were included in this study.

NACC: The National Alzheimer's Coordinating Center is a multicenter data repository that integrates data collected across Alzheimer's Disease Research Centers in the United States. The MRIs were acquired using non-harmonized protocols across centers, resulting in substantial heterogeneity in scanners, acquisition protocols, and image resolutions. To ensure sufficient image quality and spatial resolution for the current analysis, we excluded 36 T1w MRI scans with a voxel size greater than 2 mm in any dimension. After exclusion, a total of 2,920 T1w MRI scans from NACC were included in this study.

NIFD/FLDNI: The Frontotemporal Lobar Degeneration Neuroimaging Initiative study focuses on characterizing the progression of different subtypes of frontotemporal dementia (FTD). NIFD used a shared acquisition protocol across participating imaging centers. A total of 338 T1-weighted (T1w) MRI scans from NIFD were included in this study.

CCNA: The Canadian Consortium on Neurodegeneration in Aging was designed to investigate multiple causes and forms of neurodegeneration rather than focusing on a single neurodegenerative disorder. CCNA is a multisite and multivendor cohort, with MRI data acquired across different scanners using a harmonized acquisition protocol.

CIMA-Q:The Consortium for the Early Identification of Alzheimer's Disease–Quebec focuses on identifying early biomarkers and characterizing pathological changes across the Alzheimer's disease-related cognitive spectrum. CIMA-Q is a multicenter longitudinal cohort in Quebec with harmonized MRI acquisition protocols across imaging centers.

ALLFTD: The ARTFL-LEFFTDS Longitudinal Frontotemporal Lobar Degeneration study focuses on both sporadic and familial forms of FTD. Similar to NIFD, MRI data were acquired across multiple imaging sites using harmonized acquisition protocols.

PrevenAD: The PResymptomatic EValuation of Experimental or Novel Treatments for Alzheimer's Disease study longitudinally follows participants who were cognitively unimpaired at enrollment but had a first-degree family history of sporadic AD-like dementia. T1-weighted (T1w) MRI scans were acquired at a single imaging site.

HCP Aging: The Human Connectome Project Aging focuses on characterizing changes in brain structure and connectivity across healthy aging. MRI data were acquired across four imaging sites using harmonized acquisition protocols and matched 3T Siemens scanners. High-resolution T1-weighted (T1w) images were acquired at 0.8-mm isotropic resolution.

HCP Adults: The Human Connectome Project Young Adult dataset consists primarily of healthy young adults aged 22–35 years. MRI data were acquired at a single imaging site at Washington University

CamCAN: The Cambridge Centre for Ageing and Neuroscience is a cross-sectional study primarily designed to investigate age-related changes across the adult lifespan. The released T1w MRI images from CamCAN are deidentified. To mitigate potential bias associated with the defacing techniques used in CamCAN, we manually created defacing masks for the CamCAN images and transformed these masks into ICBM standard space. A total of 650 T1w MRI scans from CamCAN were included in this study.

### S2 Subsets of datasets used for hyper-parameter selection

#### Dataset A (QC-passed scans only)

Dataset A was used for the template-input ablation experiment and hyperparameter tuning. As shown in Table S1, 5,000 MRI scans from five datasets were used for training, while the remaining scans from these datasets were assigned to the validation and test sets. CamCAN, PREVENT-AD, and NIFD were additionally used as independent external test datasets. The training, validation, and test sets were stratified to preserve the distributions of image resolution and scanner characteristics across the three splits. Scans with unique scanner–resolution combinations that could not be stratified across the three sets were assigned to the training set.

**Table S1**. Detailed information on Dataset A used for template-input ablation experiment and penalty factor selection, including clinical cohorts, and sample sizes for the training, validation, and test sets.

| Dataset | Clinical Cohorts / Populations | Train | Validation | Test |
|---|---|---|---|---|
| ADNI | Cognitively unimpaired, MCI, AD | 1000 | 431 | 432 |
| ALLFTD | FTD | 1000 | 394 | 395 |
| NACC | Cognitively unimpaired, MCI, AD | 1000 | 814 | 814 |
| PPMI | Healthy controls, PD | 1000 | 814 | 140 |
| UKBB | Population-based aging cohort | 1000 | 1000 | 1000 |
| CamCAN | Healthy Aging | - | - | 610 |
| NIFD | FTD | - | - | 336 |
| PreventAD | Cognitively unimpaired individuals with a family history of AD | - | - | 295 |

### Dataset B (QC passed scans only)

Dataset B was used to evaluate the effect of training sample size on model performance. As shown in Table S2, five datasets were held out as independent test datasets. From the remaining datasets, 20% of the scans were reserved for validation, while the remaining scans formed the candidate training pool. Training sets of different sizes were then proportionally sampled from this pool, ranging from 4,000 to 15,000 scans in increments of 1,000. Five-fold cross-validation was performed for each training sample size to evaluate the variability in model performance.

**Table S2. Composition of Dataset B used for the training sample size analysis, including the numbers of scans assigned to the training pool, validation set, and test datasets.**

| Datasets | Sample size for Training Pool | Sample size for validation | Sample size for test |
|---|---|---|---|
| UKBB | 7600 | 1900 | - |
| ADNI | 234 | 58 | - |
| PPMI | 1874 | 468 | - |
| NACC | 2078 | 520 | - |
| CCNA | 585 | 146 | - |
| ALLFTD | 1425 | 356 | - |
| HCP Adults | 834 | 208 | - |

| | | | |
|---|---|---|---|
| HCP Aging | 507 | 127 | - |
| CIMA-Q | - | - | 254 |
| CamCAN | - | - | 610 |
| PrevenAD | - | - | 275 |
| HABS-Aging | - | - | 253 |
| NIFD | - | - | 339 |

### Dataset C (Real passed and real failed scans from ANTs)

Dataset D was used to train and evaluate the binary classification module. All MRI scans that failed manual quality control were included. An equal number of quality-control-passed scans were then selected to match the failed scans based on image resolution, field strength, and scanner information. This procedure produced a balanced dataset with similar numbers of QC-passed and QC-failed scans. Most of the QC-failed scans were from UK Biobank.

**Table S3. Composition of Dataset C used for training and evaluating the binary classification module, including the numbers of QC-passed and QC-failed MRI scans from each dataset.**

| Datasets | Sample size for Training set | Sample size for validation set | Sample size for test set |
|---|---|---|---|
| UKBB | 3674 | 919 | 1149 |
| PPMI | 470 | 111 | 142 |
| NACC | 210 | 47 | 65 |
| CCNA | 97 | 24 | 31 |
| ADNI | 61 | 15 | 20 |
| PreventAD | 51 | 13 | 16 |
| ALLFTD | 36 | 9 | 11 |
| CamCAN | 26 | 6 | 8 |
| CIMAQ | 18 | 4 | 6 |
| HABS Aging | 13 | 3 | 4 |
| HCP Aging | 10 | 3 | 3 |
| HCP Adults | 8 | 2 | 2 |

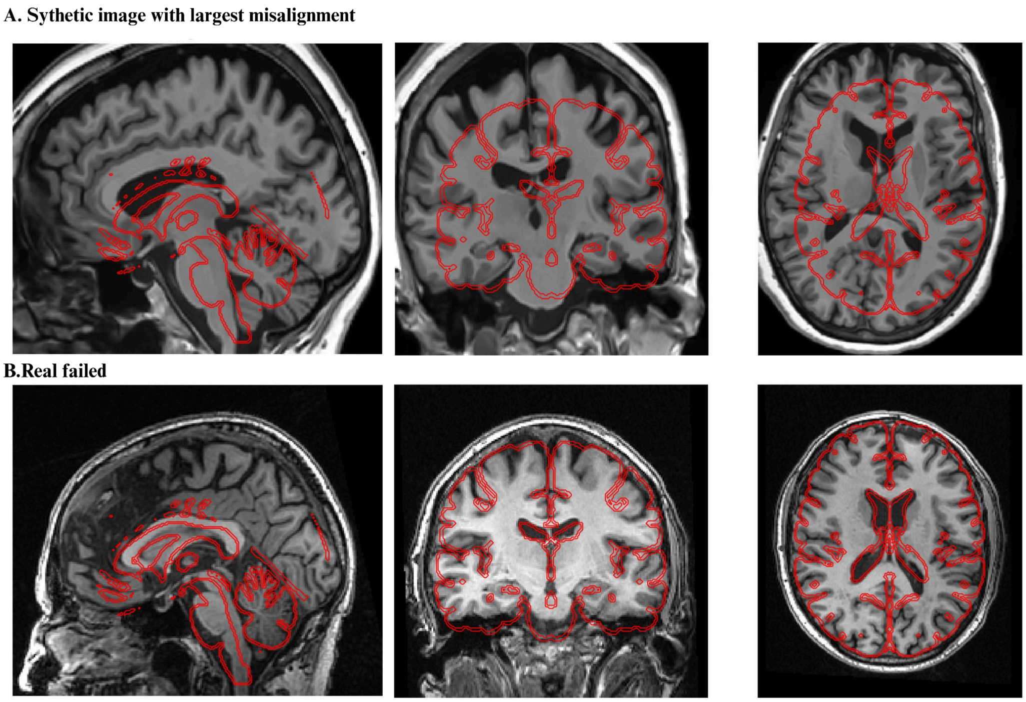


**Figure S1. Representative middle axial, coronal, and sagittal slices of real QC-failed and synthetically misaligned scans with the largest misalignment**

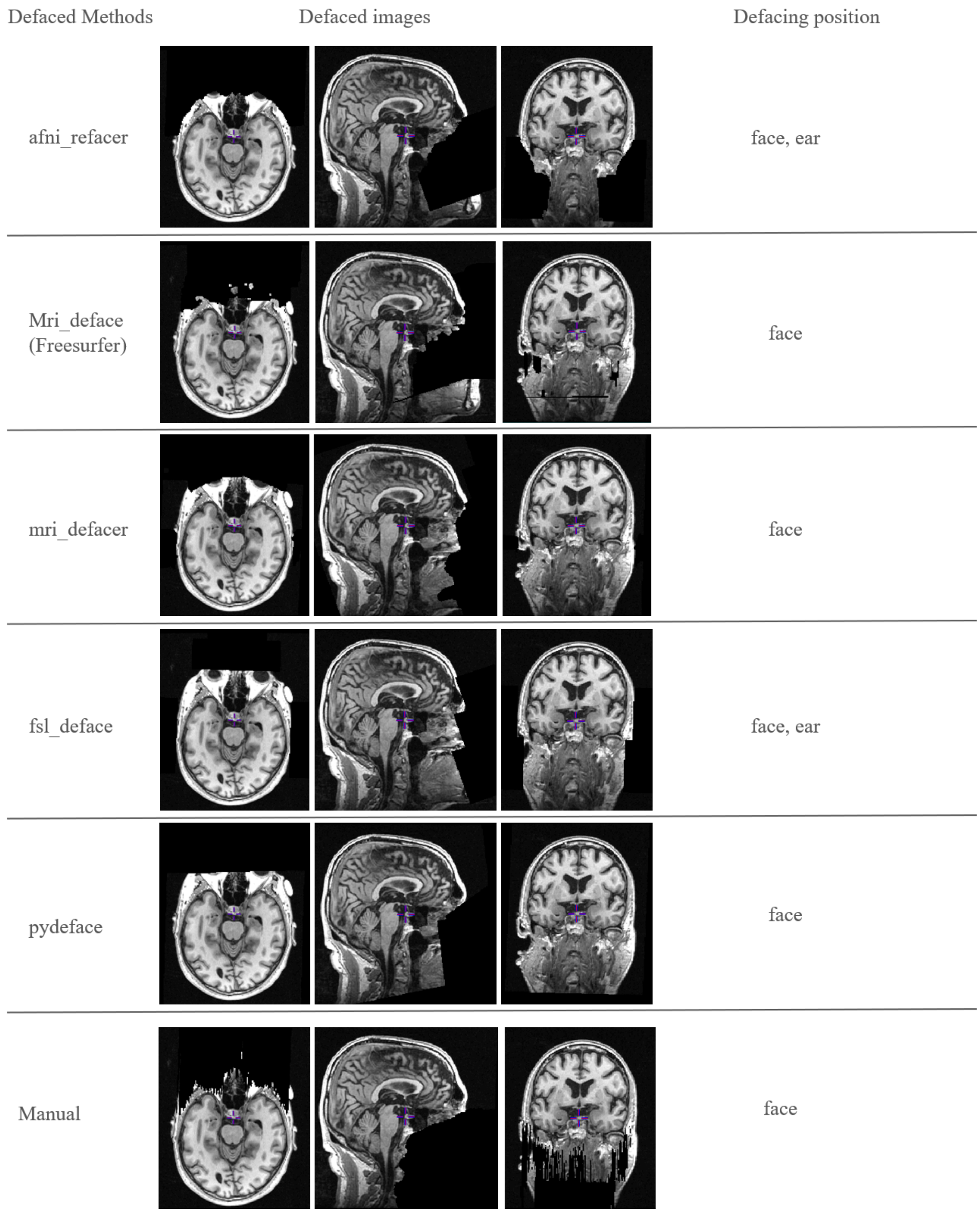


**Figure S2. Examples of different facial masks and defaced MRI scans**

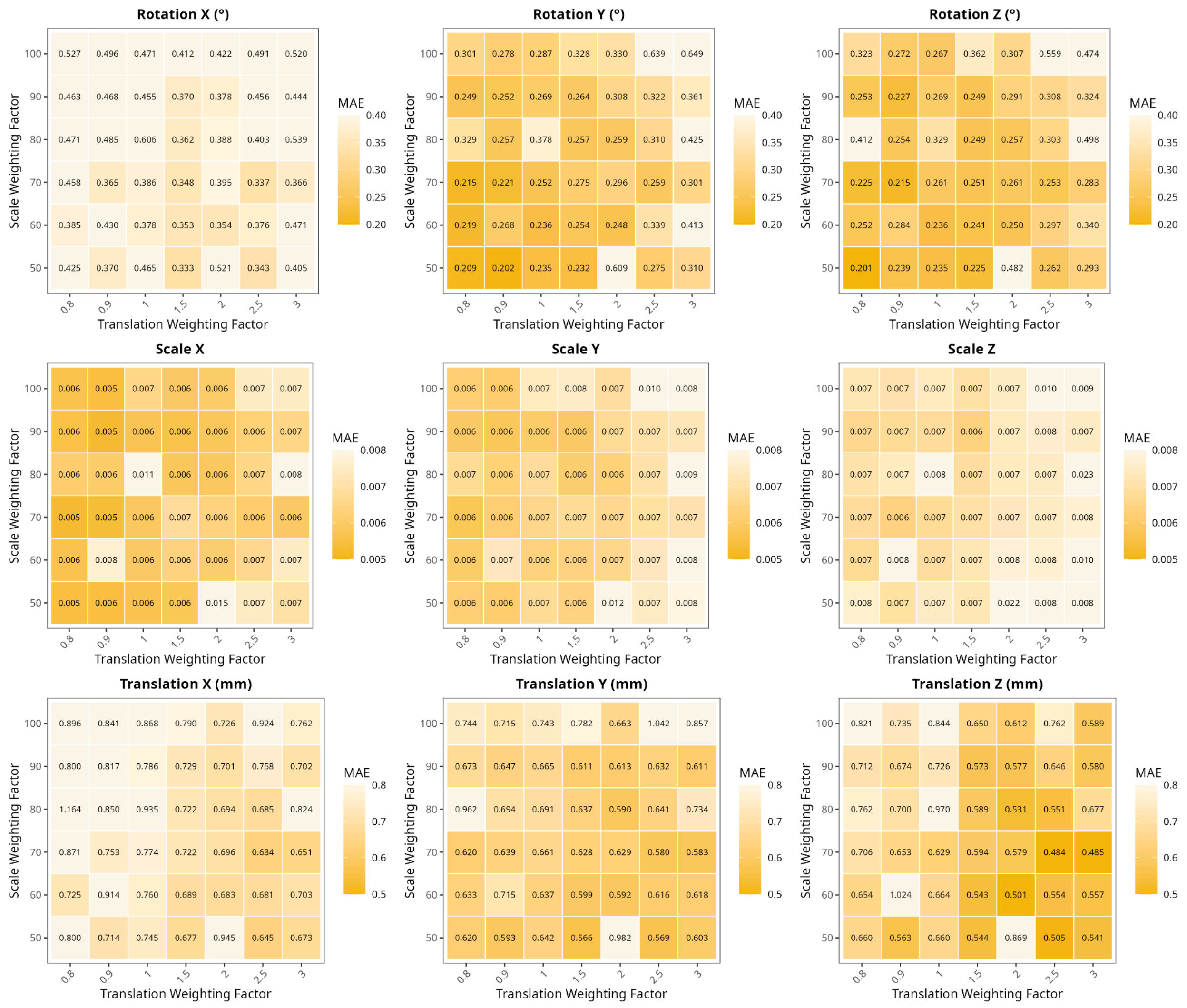

**Figure S3. RMSE of Grid search heatmaps for each affine parameter in test set**

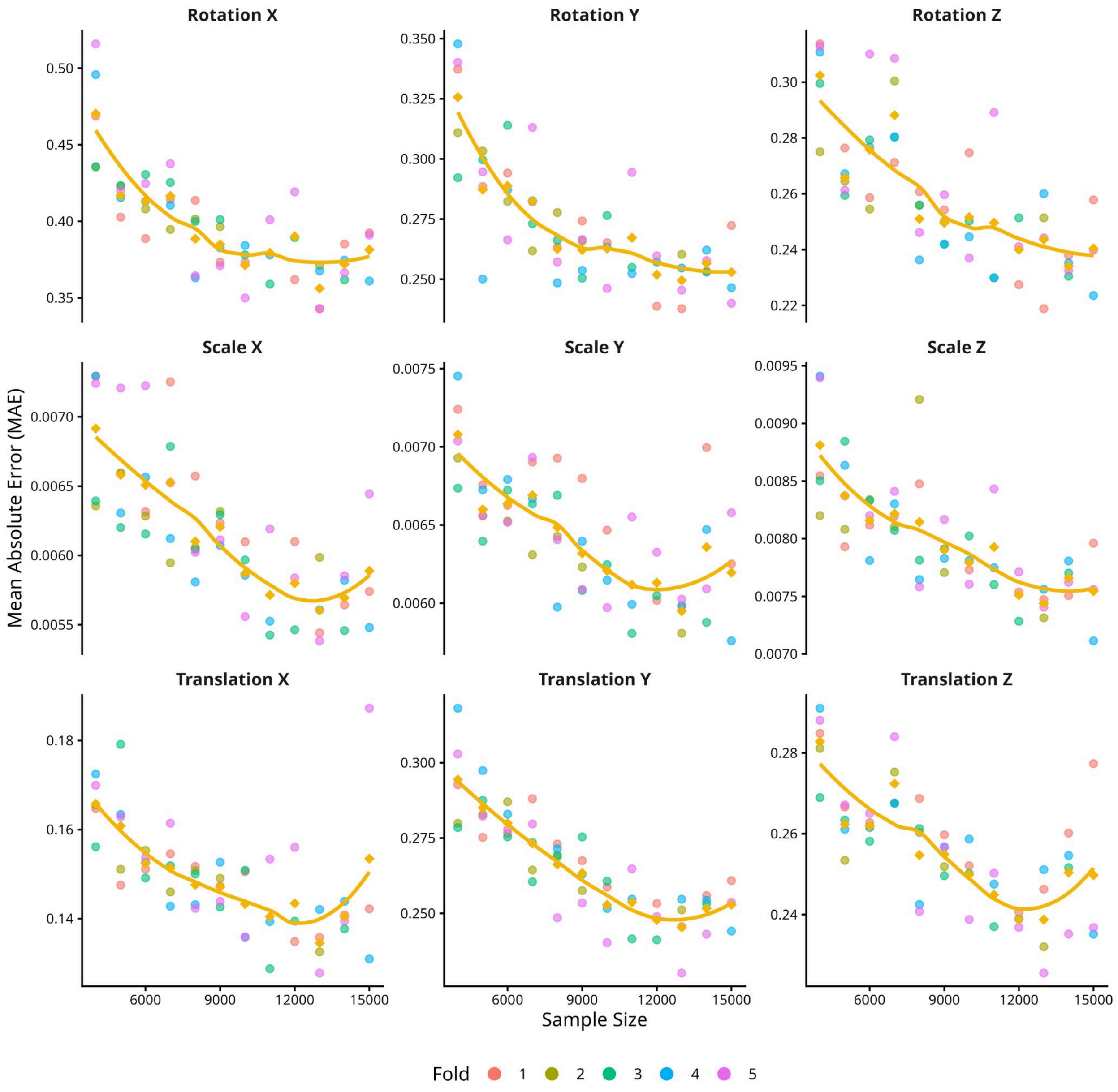


**Fig S4. MAE of the affine parameters  for the residual parameter prediction models trained with different sample sizes on the independent test sets** . Colored points represent the mean MAE across all the MRI scans for each of the five folds, and the yellow line represents the mean RMSE across the five folds at each training sample size.

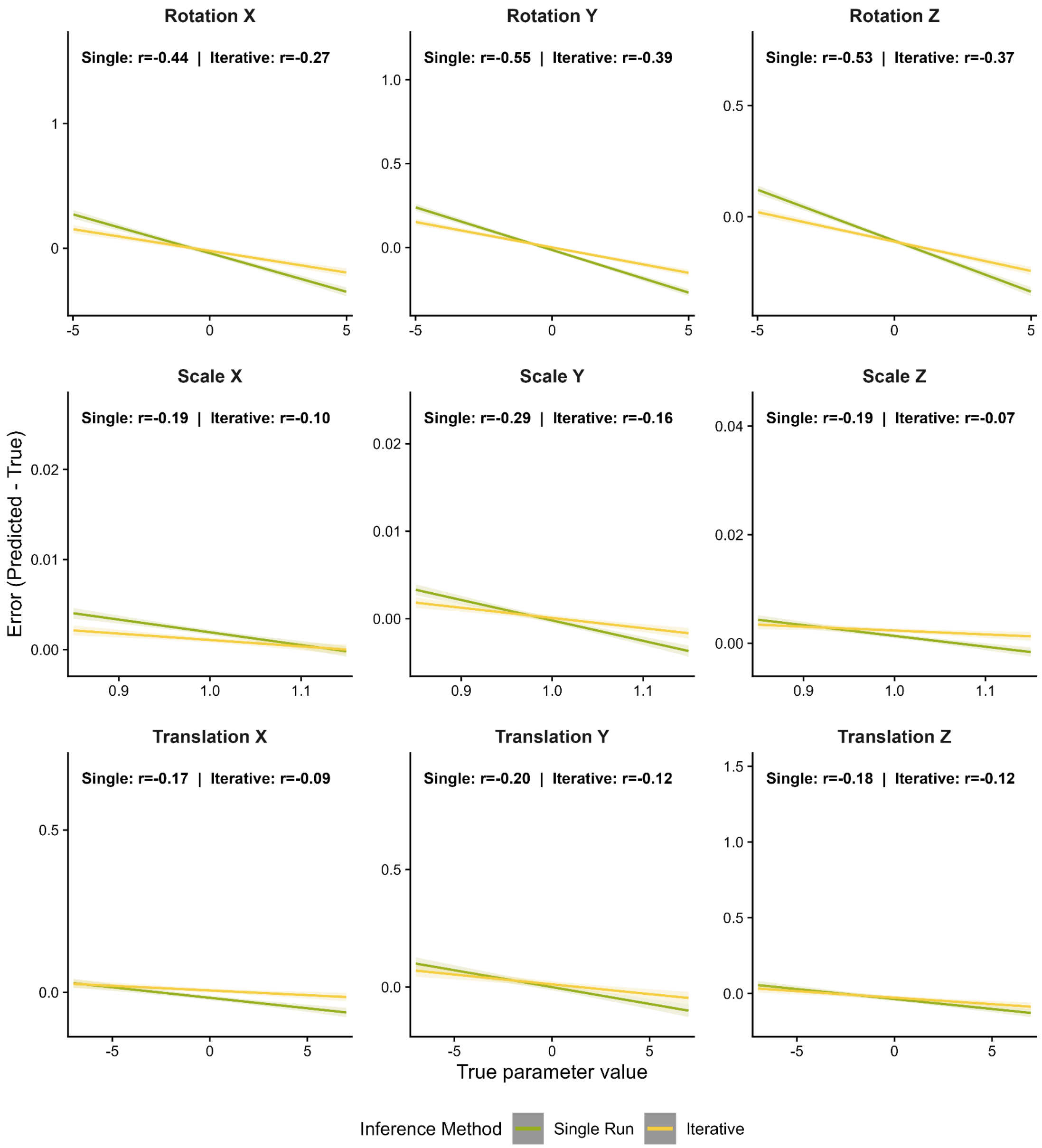


Fig S5. Correlation between prediction errors and ground-truth affine parameters for different inference strategies. Single-stage and iterative inference are shown in green and yellow, respectively. The correlation coefficients between the prediction errors and ground-truth values for each affine parameter are shown at the top of each subfigure.

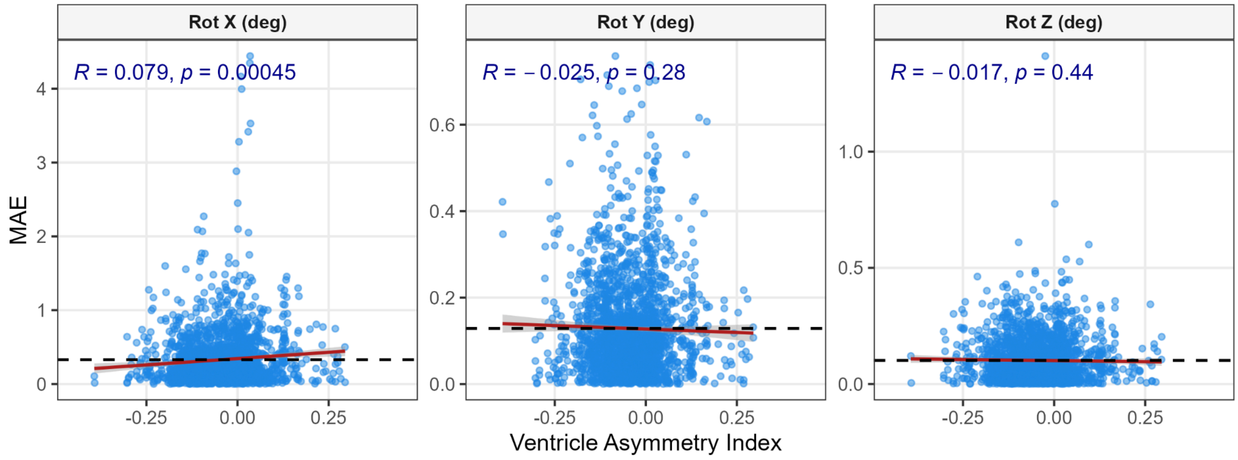


**Fig S6. Correlation between the MAE of rotation parameters in real QC-passed ADNI scans and the ventricular asymmetry index.** The ventricular asymmetry index was calculated as the difference between the summed DBM values in the left and right ventricles divided by their sum: (left − right)/(left + right).

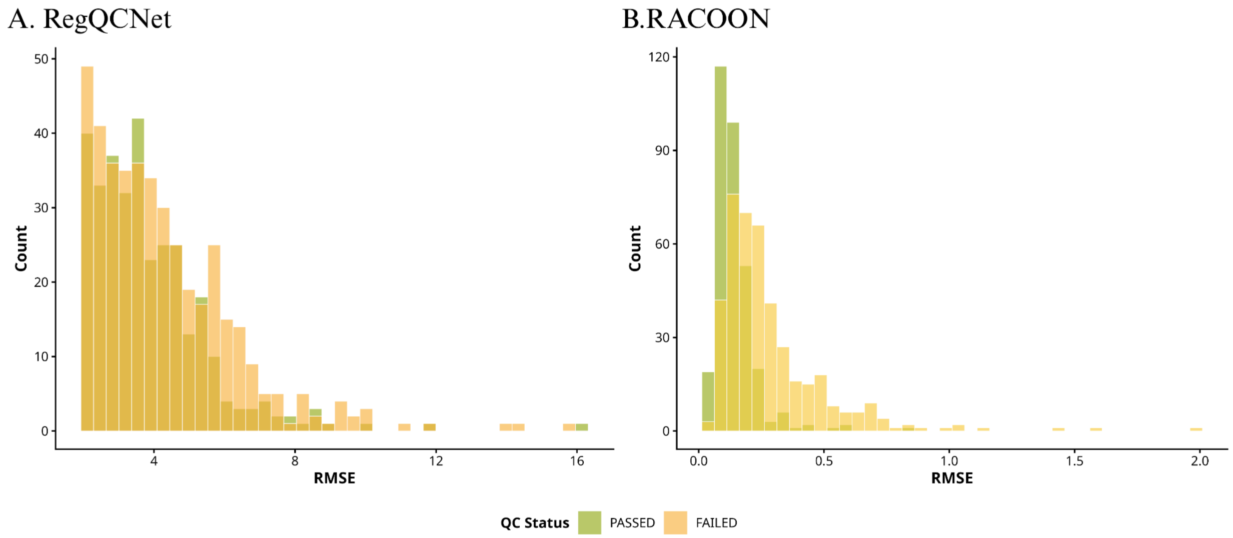


**Fig S7. Distributions of RMSE predicted by RegQCNET (A) and RACOON (B) for QC-passed and QC-failed MRI scans in the test sets.**

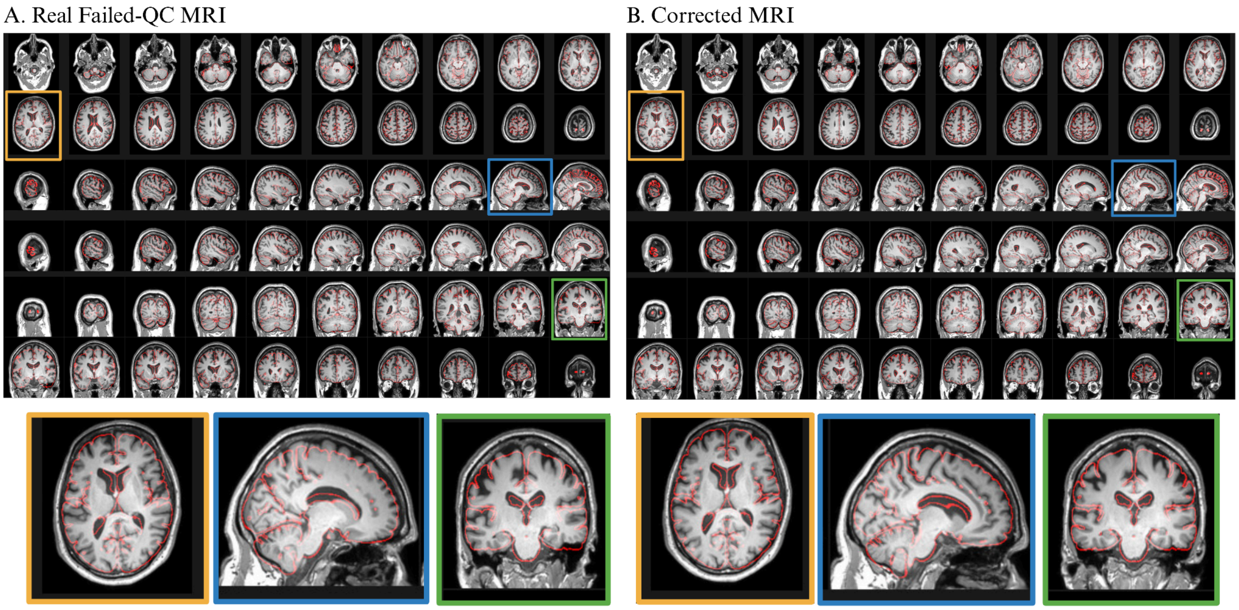


**Fig S8. Example of a QC-failed MRI scan (A) and the corresponding MRI scan corrected by RACOON (B).** For QC-failed MRI scan, translation errors are visible in the sagittal view, rotation errors in the axial and coronal views, and a shearing error in the axial view. RACOON corrected the translation and rotation errors and improved the overall alignment. However, the shearing error remains visible in the axial view because shearing parameters are not included in the current correction module.
